# Development and Validation of a Novel LC-MS/MS Based Proteomics Method for Quantitation of Retinol Binding Protein 4 (RBP4) and Transthyretin (TTR)

**DOI:** 10.1101/2025.02.27.640583

**Authors:** Aprajita S. Yadav, Katya B. Rubinow, Alex Zelter, Aurora K. Authement, Bryan R. Kestenbaum, John K. Amory, Nina Isoherranen

## Abstract

Retinol binding protein 4 (RBP4), the circulating carrier of retinol, complexes with transthyretin (TTR) and is a potential biomarker of cardiometabolic disease. However, RBP4 quantitation relies on immunoassays and western blots without retinol and TTR measurement. A liquid chromatography-tandem mass spectrometry (LC-MS/MS) method for simultaneous absolute quantitation of circulating RBP4 and TTR is critical to establishing their biomarker potential. Surrogate peptides with reproducible, linear LC-MS/MS response were selected. Purified proteins were used as quantitation standards and heavy-labelled peptides as internal standards. Matrix effects were evaluated. The validated method was applied to measure inter- and intra-individual variability in RBP4 and TTR concentrations in healthy individuals and patients with diabetic kidney disease. Quantitation was linear for the clinically relevant concentration ranges of RBP4 (0.5-6 µM) and TTR (5.8-69 µM). Assay inter-day variability was <12% and precision within 5%. The inter-individual variability for RBP4 and TTR concentrations was 18-26%, while intra-individual variability was similar to assay variability. RBP4 and TTR quantitation correlated with commercially available ELISA assays. The developed LC-MS/MS method enables simultaneous absolute quantitation of RBP4 and TTR in serum and plasma that can be applied to clinical biomarker studies and stoichiometric measurements of circulating RBP4, TTR, and retinol.

## INTRODUCTION

Retinol binding protein 4 (RBP4) is the blood carrier of retinol (vitamin A). RBP4 is a 21 kDa protein and is filtered by the kidney.^1,2^ RBP4 forms a complex with transthyretin (TTR).^1^ Formation of the complex prevents glomerular filtration and increases the half-life of RBP4.^3^ TTR is a 55 kDa homotetramer that is stabilized by RBP4 binding.^4,5^ In healthy individuals, in the absence of nutritional vitamin A deficiency, the plasma concentrations of retinol^6^, RBP4^2^, and TTR^5^ have little variability between individuals.^7^ Blood RBP4 concentrations are, however, affected by dietary vitamin A status^8,9^ and by disease.^4,10,11^ For example, with decreased kidney function, plasma RBP4 concentrations are increased suggesting that RBP4 may be a biomarker of kidney disease.^12^ Similarly, increased concentrations of RBP4 have been proposed as a biomarker of diabetes, cardiovascular disease,^13,14^ preeclampsia and gestational diabetes,^15,16^ although during pregnancy, RBP4 concentrations appear to decrease.^17^ RBP4 is typically measured in isolation, and TTR concentrations along with the formation of RBP4-TTR complex are not considered in the context of RBP4-disease associations, despite the biological relevance.^11^ The use of RBP4 and TTR as biomarkers individually or simultaneously is hindered by the lack of validated methods to quantify RBP4 and the limited data on intra-individual variability of RBP4 and TTR plasma concentrations.

RBP4 and TTR concentrations have typically been measured by western blots or enzyme linked immunoassays (ELISA).^5,11^ Intact protein mass spectrometry has also been used to differentiate misfolded and correctly folded TTR.^18,19^ The ELISA and western blot methods have shortcomings such as limited dynamic range, variability between assays, and interference from disease states, matrix effects, or anticoagulants.^20,21^ These shortcomings in RBP4 measurement methodology in particular have been suggested to result in inconsistent clinical findings regarding associations of RBP4 concentrations and disease progression.^11,21^ Whether the RBP4-TTR complex formation interferes with antibody-based assays is currently unknown. Liquid chromatography-tandem mass spectrometry (LC-MS/MS) based protein quantitation methods hold promise as an alternative strategy to immunoassays and western blots. LC-MS/MS based methods can advance the understanding of RBP4 and TTR as biomarkers or etiologic factors in metabolic and chronic diseases.

The goal of this study was to develop an LC-MS/MS based assay to quantify RBP4 and TTR in human plasma and serum. A robust and sensitive method with stable isotope labeled (SIL) peptides as internal standards was developed and validated. Matrix effects were evaluated, and the RBP4 and TTR quantitation by the LC-MS/MS method was compared to the ELISA based quantitation using a set of samples from patients with diabetic chronic kidney disease. We also established inter- and intra-individual biological variability of RBP4 and TTR serum and plasma concentrations by analyzing multiple matched serum and plasma samples from healthy individuals.

## MATERIALS AND METHODS

### Chemicals, Reagents, and Reference Materials

Sodium deoxycholate, iodoacetamide, ammonium bicarbonate, human serum albumin (PN A9511), bovine serum albumin (PN A6003), trifluoroacetic acid, and yeast enolase were purchased from Sigma-Aldrich (St. Louis, MO). Previously collected serum from healthy human volunteer was used for assay development.^22^ Dithiothreitol (DTT), tris(2-carboxyethyl)phosphine (TCEP) solution, and Ringer’s solution were purchased from Thermo Fisher Scientific (Waltham, MA). Trypsin Platinum (PN VA9000) was purchased from Promega (Madison, WI). RapiGest SF was purchased from Waters (Milford, MA). Optima LC-MS grade acetonitrile, water, acetic acid, and formic acid were purchased from Fisher Scientific (Pittsburg, PA). Phosphate buffered saline (PBS) was purchased from Corning (Corning, NY).

Purified RBP4 was purchased from multiple sources (**SI Table S1**). TTR purified from human serum was purchased from Bio-Rad Laboratories and was >96% pure per vendor certificate of analysis. The purity of purchased proteins was further confirmed by gel electrophoresis and Coomassie staining. Lyophilized RBP4 and TTR were reconstituted according to supplier guidelines to 1 mg/mL using PBS (47.46 µM RBP4 and 72.67 µM TTR). Protein concentrations were measured from absorbance at 280 nm with a Thermo Nanodrop using an extinction coefficient (ε) of 34,295 M^-1^cm^-1^ for RBP4 and 18,450 M^-1^cm^-1^ for TTR (Expasy ProtParam^23^). The protein concentrations of Bio-Rad RBP4 and TTR (>96% pure per vendor certificate of analysis) were confirmed by bicinchoninic acid (BCA) assay (Thermo Scientific). Based on the BCA assay, the accuracy across three concentrations of protein (0.3, 0.5, and 0.7 mg/mL) was 0.5-2.5% for RBP4 and 4.6-6.3% for TTR. The concentrations of the final reference solutions of RBP4 and TTR were confirmed using amino acid analysis as previously described.^24^ Amino acid analysis (**SI Table S2**) resulted in RBP4 and TTR protein concentrations within 1.4% and 6.9% of their nominal values, respectively, confirming reference solution concentrations.

Stable isotope labeled (SIL) peptides FSGTWAYAMAK[^13^C ^15^N], YWGVASF[^13^C_9_ ^15^N]LQK, and GSPAINVAVHVFR[^13^C ^15^N] were purchased from Thermo Fisher Scientific (Waltham, MA) and were >95% pure.

### Identification of surrogate peptides and digestion time course

The amino acid sequences for human RBP4 and TTR in FASTA format were retrieved from UniProtKB database (P02753 and P02766, respectively). RBP4 and TTR were digested *in silico* using Skyline v22^25^ (Seattle, WA) to identify tryptic peptides. Resulting tryptic peptides with 7-25 amino acids and predicted m/z range of 50 – 1500 were considered further. Nonspecific peptides in human plasma were screened as previously described^26^ and excluded. For the remaining peptides, precursor ions with 2 or 3 charges, peptide fragmentation (b and y ions), and instrument and peptide specific optimal declustering potential and collision energies were predicted in Skyline to incorporate into initial method development.

To test peptide detection and fragmentation, purified recombinant RBP4 and TTR (diluted to 400 nM using 100 mM ammonium bicarbonate pH 7.8) were digested as previously described.^26^ In brief, samples were incubated with DTT followed by addition of deoxycholate before samples were heated to 95°C to denature proteins. Subsequently iodoacetamide was added to samples before trypsin digestion. The potential tryptic peptides identified in silico (n=8 for RBP4 and n=5 for TTR) were monitored in the digested samples using multiple MS/MS transitions to determine which peptides and transitions were detectable. The details of the monitored and detected peptides and MS/MS transitions are included in **SI Table S3**. The LC method was as described below for LC-MS/MS method.

The digestion efficiency was evaluated in experiments of trypsin digestion time course (3, 5, 15, and 20 h digestion time) using previously reported general digestion conditions.^26^ In these experiments all peptides detected from purified RBP4 (n=7) and TTR (n=3) in the initial experiments were monitored together with the corresponding peptides with a single missed cleavage if the peptide contained a ragged end (**SI Table S3**). The trypsin digestion time course was evaluated on three different days. Each day digestion time course was evaluated in triplicate digestions of purified TTR (diluted to 200 nM using 100 mM ammonium bicarbonate pH 7.8), purified RBP4 from four vendors (diluted to 20 nM using 100 mM ammonium bicarbonate pH 7.8), and serum from a healthy human volunteer diluted 100-fold with 100 mM ammonium bicarbonate pH 7.8.

Possible methionine oxidation during sample preparation on the RBP4 peptide FSGTWAYAMAK was assessed using similar digestion conditions as described below for final digestion protocol. Eight MS/MS transitions for y and b-ions of the oxidized peptide, FSGTYAM(Oxi)AK were added to the method. The dwell time was 30 msec for each transition, and the transitions were: precursor^+2^ 589.3 > 1030.5 (y9), 943.4 (y8), 785.4 (y6), 599.3 (y5), 436.2 (y4), 365.2 (y3), 393.2 (y6^+2^), and 235.1 (b2). Peak areas from these transitions were compared to the peak areas of the unoxidized peptide in an analysis of a digested diluted RBP4 standard (20 nM) corresponding to 2 μM RBP4 undiluted, similar to physiological circulating concentrations.

### Impact of denaturation and detergent on digestion

The impact of reducing agent and detergent on digestion efficiency and peptide detection was assessed using purified RBP4 (20 nM in ammonium bicarbonate) and TTR (200 nM in 100 mM ammonium bicarbonate) and human serum diluted 100-fold with ammonium bicarbonate. Reduction with DTT or TCEP was compared following digestion of the samples in the presence of either 8 μL of fresh 100 mM DTT or 50 mM TCEP dissolved in 100 mM ammonium bicarbonate added to 40 µL samples. Remaining protocol including denaturation, alkylation, 5 h trypsin digestion, and quenching were kept the same between digestions. The peak area ratios of RBP4 and TTR peptides (FSGTWAYAMAK, and GSPAINVAVHVFR) to their corresponding labeled peptides were compared between the digestions with different reducing agents.

The effect of a surfactant or denaturation on digestion efficiency was also assessed using DTT as the reducing agent. Samples were prepared in triplicate and the effects of detergent (deoxycholate versus Rapigest), and temperature (25°C, 60°C or 95°C) were tested. RapiGest powder was reconstituted in 100 mM ammonium bicarbonate pH 7.8, and the final concentration of RapiGest in the digestion was 0.1%. The reduction, alkylation, trypsin digestion for 5 hours and quenching the digestion with acetonitrile containing 8% TFA and internal standard remained as described above. RapiGest incubations are recommended at 60°C and were conducted at 60°C or 95°C to compare conditions with only one variable altered at a time between detergents. The RBP4 and TTR peptide (FSGTWAYAMAK, and GSPAINVAVHVFR) peak areas were divided by the SIL peptides to obtain peak area ratios.

### LC-MS/MS method

Peptides were separated using an Aeris Peptide column (50 x 2.1 mm, 1.7 µm) with a SecurityGuard Ultra C18-peptide cartridge (Phenomenex, Torrance, CA) on an Agilent 1290 LC (Agilent, Santa Clara, CA).^26^ Injection volume was 10 µL for RBP4 analysis and 3 µL for TTR analysis. The column oven was set to 40°C. The mobile phases were water (A) and acetonitrile (B), both with 0.1% formic acid, at flow rate of 0.4 mL/min. Gradient elution was from 3% B for 3.5 min, increased linearly to 40% B over 8.5 min, increased to 95% B over 0.5 min, and held at 95% B for 2.5 min before returning to starting conditions.

Peptides were detected using a SCIEX 5500 QTRAP MS (Sciex, Framingham, MA). Isotope labelled peptides were infused to optimize mass spectrometer parameters. The final, optimized, mass spectrometer parameters for the surrogate peptides used for quantification are summarized in **SI Table S4**. Three MS/MS transitions were selected for each quantitation peptide and summed peak area was used.

### Final Digestion Protocol

For sample analysis and method validation, human serum samples were diluted 100-fold with 100 mM ammonium bicarbonate pH 7.8. For sample preparation 40 µL of diluted samples, standard curves and quality control (QC) samples were aliquoted into 96-well PCR plates with 20 µL of 100 mM ammonium bicarbonate (pH 7.8), 8 µL 100 mM dithiothreitol (DTT), and 2 µL of 8 ng/µL yeast enolase as a process control. Samples were incubated for 20 min at room temperature to reduce disulfide bonds before addition of 10 µL of 10% sodium deoxycholate in 100 mM ammonium bicarbonate. Proteins were denatured by incubation at 95°C for 10 min in a ThermoMixer. Under yellow light, 8 µL of 200 mM iodoacetamide was added, and the samples were incubated for 20 min at room temperature. Samples were digested with trypsin at 37°C for 5 h at 400 rpm in a ThermoMixer. Digestions were quenched with 40 µL of acetonitrile with 8% TFA containing internal standard peptides (50 nM FSGTWYAMAK[^13^C ^15^N] and YWGVASF[^13^C_9_ ^15^N]LQK, and 250 nM GSPAINVAVHVFR[^13^C ^15^N]). The plate was centrifuged at 3,000 *g* for 30 min at 4°C, the orientation of the plate flipped 180° and centrifuged for another 30 min, and supernatants transferred to a new 96-well plate for LC-MS/MS analysis.

### Method validation

Validation was performed according to Food and Drug Administration guidance^27^, including assessment of linearity, parallelism, matrix effects, lower limit of quantification (LLOQ), precision, accuracy, stability of digested peptides, and carryover. Enolase peptides (**SI Table S4**) were monitored as a process control.^28^

Signal linearity at concentrations relevant for analysis was first confirmed by serial dilution of digested purified protein diluted with ammonium bicarbonate: acetonitrile (7:3) to the concentration corresponding to diluted samples, 60 nM and 40 nM of RBP4 done as parallel serial dilutions and 800 nM for TTR. Parallelism was assessed by quantifying human serum from healthy volunteer diluted with mouse serum at 100%, 75%, 50%, 25%, and 0% human serum, with each dilution tested in triplicate. Potential bias in the slope of the dilution series was evaluated for both RBP4 and TTR measurements.

Matrix effects and assay specificity were evaluated following 5 h trypsin digestion of RBP4 (20 nM) and TTR (200 nM) spiked into different surrogate matrices including human serum albumin (HSA) or bovine serum albumin (BSA) (0.6 mg/mL) in 1% Ringer’s in 100 mM ammonium bicarbonate pH 7.8, fetal bovine serum (FBS), mouse serum, and mouse plasma. The peak area ratio of the selected surrogate peptides for RBP4 and TTR quantification to their isotope labeled internal standard peptides was quantified.

Instrument variance, including autosampler and detection variability, was assessed by calculating coefficient of variance (%CV) of the summed MS/MS transition peak area for the peptide of interest from twelve replicate injections.

Standard curves and QCs were prepared using the concentration verified RBP4 and TTR (Bio-Rad) solutions in PBS. Diluted stock solution of 600 nM RBP4 and 6.9 µM TTR was prepared by adding 8.1 µL of 1 mg/mL RBP4 and 61 µL of 1 mg/mL TTR to PBS for a total volume of 640 µL. Separate intermediate stocks were used to generate standard curves and QCs. Standard curves were then prepared by spiking appropriate volumes of the diluted stock to the surrogate matrix of 0.6 mg/mL BSA in 1% Ringer’s in 100 mM ammonium bicarbonate. The standard curves included 8 non-zero points corresponding to 0.5 – 6 µM RBP4 and 5.8 – 69.3 µM TTR in plasma and diluted 100-fold with ammonium bicarbonate for analysis. QC samples were prepared at four concentrations: lower limit of quantitation (LLOQ), low QC (LQC), medium QC (MQC), and high QC (HQC) corresponding to concentrations of 0.6, 0.9, 1.8, 3.6 µM for RBP4 and 6.9, 10.4, 20.8, and 41.6 µM for TTR with a 100-fold dilution. The LLOQ was determined per the FDA Bioanalytical Guidance as the lowest verified concentration with accuracy and precision within 20%.^27^ Due to the expected circulating concentration ranges of RBP4 and TTR, a lower LLOQ was not pursued despite the signal to noise being considerably over 10 at the defined LLOQ. The reported LLOQ does not reflect the achievable lower limit of quantification for this method.

A minimum of three QC samples per day were digested and analyzed across nine days to determine inter-day variance. In addition, a pooled mixture of serum from ten individuals was prepared and aliquoted for analysis as a pooled QC. The pooled QC was run in triplicate with every run and intra- and inter-day variance were calculated for the pooled QC as well. Intra-day variance was calculated from four QC samples per concentration within a day and the mean intra-day variance from nine days is reported. Accuracy (% error or bias) was calculated as (nominal – observed)/nominal. The observed concentration was the concentrations measured on that day for the QC sample based on the standard curve.

Freeze-thaw and autosampler stability of digested peptides was determined by quantifying QC samples that were either frozen at – 20°C for two freeze-thaw cycles or kept at 4°C for 24 h, respectively.

The internal standard SIL peptides do not need digestion to be detected in comparison to the sample surrogate peptides. Hence they do not necessarily behave the same as purified protein through sample preparation. The SIL peptides were added to the samples after digestion to control for signal stability in the analysis and enolase peptides were monitored as a control for digestion efficiency. To confirm lack of bias due to addition of SIL peptides after digestion, the effect of adding SIL peptides at the beginning of sample processing compared to at the end of digestion was evaluated. Parallel digestions of standard curves, three sets of QC samples, and pooled QCs were conducted in triplicate. In one set, samples were processed as described above and SIL peptides were added with quenching of the digestion. In the second set, SIL peptides were added to the samples before digestion (together with DTT and enolase). For this set, digestions were quenched with the addition of acetonitrile containing 8% TFA and no SIL peptides. Percent error was calculated at each QC concentration and for the pooled QC in both sample sets. The % difference between the methods of SIL peptide addition was calculated as the difference between the mean measured concentration when SIL peptides were added before digestion and the mean concentration when they were added after digestion, divided by the measured concentration when SIL peptides were added after digestion and multiplied by 100.

### Method comparison

A subset of human plasma samples (n=49) were obtained from the Seattle Kidney Study, a prospective cohort study of individuals with chronic kidney disease (CKD).^29^ All patients provided written informed consent, and the study protocol was approved by the Institutional Review Board at the University of Washington (STUDY00001067). The samples were from patients diagnosed with diabetes, with geometric mean (IQR, interquartile range) age of 58 (55, 70) years, estimated glomerular filtration rate of 48 (35, 64) mL/min/1.73m^2^, and 31 (63%) participants were female. These samples were analyzed by LC-MS/MS and by commercial RBP4 and TTR ELISA kits (Human RBP4 Quantikine Kit, R&D Systems, Minneapolis, MN and Human TTR ELISA Kit, Abnova, Taipei City, Taiwan) to compare RBP4 and TTR quantification by the different methods in patient samples.

There is no accepted “gold standard” reference ELISA for RBP4 and TTR, thus kits that have been previously used and reported were selected.^12,17^ Concentrations were measured according to manufacturer’s recommendations. For the ELISA assay samples were diluted 1,000-fold for RBP4 analysis and 100,000-fold for TTR analysis. Absorbance was read on a Tecan Microplate Reader (Männedorf, Switzerland) at 450 nm with a 570 nm correction for optical imperfections. Samples were quantified against a standard curve (5-point curve equivalent to 0.3-4.8 µM for RBP4 and 4-point curve equivalent to 0.9-57 µM for TTR). The two lowest RBP4 standards at 0.07 and 0.15 µM were excluded due to poor curve fit, defined as error greater than 20%. Similarly, the highest TTR standard at 227 µM was excluded from analysis due to signal saturation and suboptimal curve fit. Method concordance between the ELISA and LC-MS/MS was examined using a Bland-Altman plot, and Deming and Pearson regression models.

### Clinical variance

To evaluate intra-individual variability in RBP4 and TTR concentrations and possible differences in RBP4 and TTR quantification between plasma and serum, plasma and serum samples from a study of twelve healthy premenopausal female volunteers enrolled in a study evaluating the role of pregnancy hormones on disposition of dronabinol (ClinicalTrials.gov ID NCT04374773) were analyzed. Baseline samples were collected on two different non-consecutive days to study intra-individual variability. The study was approved by the Institutional Review Board at the University of Washington (STUDY0008064). Blood samples were collected after an overnight fast to serum separator tubes and sodium citrate tubes. Serum and plasma were separated by centrifugation at 3,000g at 4°C for 20 min and stored at −80°C until analysis.

Inter-individual variability was calculated as the geometric mean of the coefficient of variance on both baseline days separately for serum and plasma. Intra-individual variability was calculated as the mean of half the absolute difference between baseline visits for each individual divided by the mean measured concentration for all individuals.

### Data Availability

Raw data, including Skyline quantification, were deposited to the ProteomeX-change Consortium via Panorama Public^30^ and are available under the data set identifier PXD061072 at: https://panoramaweb.org/plasmaRBP4_TTR.url

## RESULTS

### Selection of surrogate peptides and optimization of digestion conditions

Seven of the eight predicted tryptic peptides for RBP4 were detected after digestion of purified protein (**Figure 1, SI Figure S1**). Only one of the detected RBP4 peptides, YWGVASFLQK (YWG), did not contain cysteine or methionine residues, or a ragged end (adjacent lysine or arginine residues). However, the YWG peptide was also detected with a missed cleavage, MKYWGVASFLQK (MKY) (**SI Figure S1D**). Of the remaining detected RBP4 peptides, FSGTWYAMAK (FSG) was considered as a second potential surrogate peptide despite the methionine present. For TTR, three of the five predicted peptides were observed (**Figure 1, SI Figure S2**), all of which contained a ragged end at either the C or N terminus (**Figure 1B**). Of these three peptides, the GSPAINVAVHVFR (GSP) peptide had the highest signal intensity (**Figure 1F**). Based on this data FSG and YWG were considered further as surrogate peptides of RBP4 and GSP of TTR.

**Figure 1.**
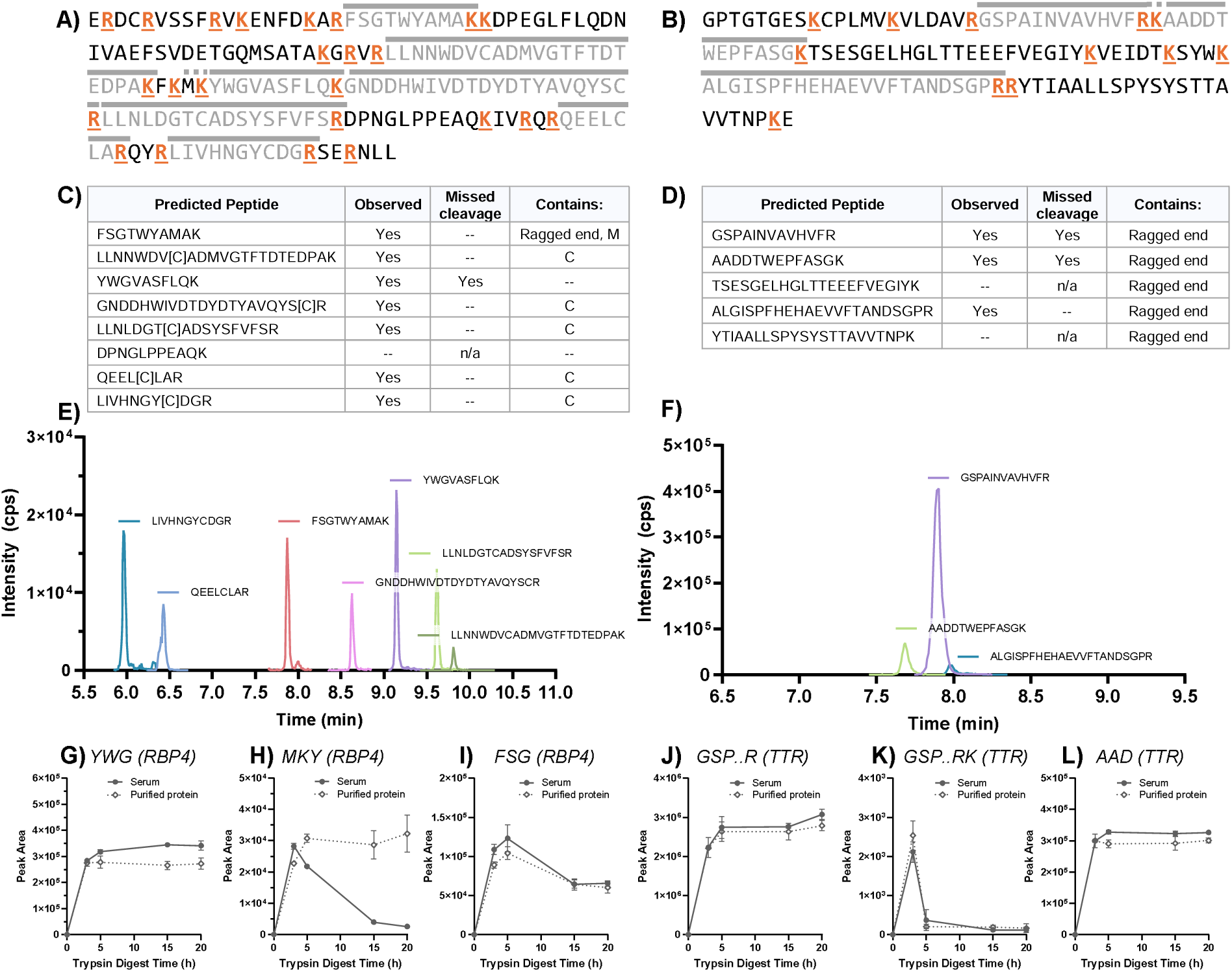
Identification and selection of RBP4 (left) and TTR (right) surrogate peptides. Amino acid sequences of RBP4 (A) and TTR (B) proteins with potential tryptic cleavage sites (lysine, K, and arginine, R) shown in orange, observed peptides noted with a gray bar above the peptide, and observed missed cleavage peptide noted with a dashed bar above the missed cleavage residues. Key characteristics of predicted peptides of RBP4 (C) and TTR (D) are summarized. Ragged end defined as KK, RK, KR, or RR. A missed cleavage was noted if a peak for the missed cleavage peptide with multiple transitions and signal-to-noise >3 was present. Representative chromatograms of observed potential RBP4 (E) and TTR (F) surrogate peptides following 5 h of trypsin digestion of purified protein are shown overlaid with the peptide sequence indicated next to each peak. The signal intensity is summed from three most abundant MS/MS transitions (**SI Table S3**). Individual observed transitions for initial peptide screen are shown in **SI Figure S1 and S2**. Trypsin digest time course of potential surrogate peptides from serum and Bio-Rad purified proteins: RBP4 peptides YWG (G), missed cleavage peptide MKY (H), and peptide FSG (I) and TTR peptides GSP (J), missed cleavage GSPAINVAVHVFRK, GSP..RK, (K), and AAD (L). The open diamonds and dotted lines show digestions of purified protein and filled circles and solid line show digestions from serum.

When purified RBP4 from different sources was digested, the peak areas and digestion time courses for YWG, MKY, and FSG peptides were not dependent on protein source (**SI Figure S3**). The presence of serum matrix in the digestion also had minimal effect on digestion time course for the majority of RBP4 and TTR peptides (**Figure 1G-L**). The only exception was the RBP4 peptide MKY with the missed cleavage for which the peak area decreased with digestion time in serum but not with purified protein (**Figure 1H**). Peak area of the missed cleavage TTR peptide GSPAINVAVHVFRK, GSP..RK, was less than 1% of the peak area of the fully cleaved peptide (GSP) and decreased substantially by 5 h digest time (**Figure 1J, K**). Based on these data, a digestion time of 5 h was considered optimal for the assay to maximize peptide signal, reduce TTR missed cleavage, and minimize peptide degradation or potential methionine oxidation. The FSG and GSP peptides were chosen as the surrogate peptides for method development.

The potential for methionine oxidation in the FSG peptide during the sample preparation process was evaluated. No distinct peak corresponding to an oxidized form of the FSG peptide was detected in a digested standard (**SI Figure S4A**). Minor peaks observed in the chromatogram had a combined peak area less than 5% of the unoxidized FSG peptide signal in the same sample (**SI Figure S4**). This suggests methionine oxidation of the FSG peptide is minimal in this method and unlikely to impact quantitation.

The choice of reducing agent (DTT or TCEP) did not affect peptide detection (data not shown) and DTT was used for further analyses. In contrast, the choice of detergent had a significant effect on digestion and peptide detection, although the effect was protein and matrix dependent. The detergent had a minor impact on FSG detection from RBP4 (**SI Figure S5**). However, deoxycholate was critical for TTR digestion and detection of the GSP peptide. The GSP peptide signal was 88 – 99% lower in the absence of deoxycholate than in its presence (**SI Figure S5**).

### Method Validation

At present, there are no reference standards available for RBP4 and TTR. Hence the concentrations of the purified protein standards were confirmed by multiple orthogonal methods which all agreed with each other. The RBP4 proteins from different sources also had similar peptide peak area response in a trypsin digestion time course (**SI Figure S3**). The peptide peak area from the digested purified protein was not significantly different (<10% difference) than the corresponding stable isotope labeled peptide peak area suggesting digestion was complete and recovery and quantification of the protein references were correct. Together with the amino acid analysis, BCA quantification and direct UV measurement, these results support accurate characterization of the reference material.

FSG and GSP peptide MS signals were linear for pre-dilution sample concentrations of 0.5 – 6 µM and 2.5 – 80 µM, respectively (**SI Figure S6A,B**). The instrument variability was 7.3% for enolase peptide AADALLLK (AAD), 7.0% for FSG, and 7.1% for GSP based on 12 replicate injections of digested pure enolase, RBP4, and TTR. When the peak area was normalized to the corresponding labeled peptides, the variability decreased to 5.2% for FSG and 3.5% for GSP.

Sample matrix can impact protein digestion and MS ionization of peptides, and some matrices can contain interfering peptides.^26,31^ No peaks were observed for the RBP4 peptide FSG in any tested matrix (**Figure 2A**), but a significant peak corresponding to TTR peptide GSP was observed in HSA in Ringer’s, suggesting that the HSA purified from human plasma is contaminated with TTR (**Figure 2B**). No contaminating signal for the FSG or GSP peptides was observed in BSA in Ringer’s, in FBS, in mouse serum or in mouse plasma. When human RBP4 and TTR were spiked into BSA in Ringer’s, mouse serum, or mouse plasma, quantified concentrations of RBP4 and TTR were within 15% of the nominal concentration at low, medium, and high QC concentrations (**Figure 2C,D**). No discernible differences were observed between the three matrices and hence BSA with 1% Ringers, mouse serum or plasma are considered acceptable surrogate matrices for the assay. Parallelism was assessed by diluting human serum with mouse serum. Measured concentrations of RBP4 and TTR across the dilution series had slopes (95% confidence intervals) of 1.05 (0.94 to 1.16) and 1.06 (0.96 to 1.15), respectively, indicating minimal matrix effect (**Figure 2E,F**).

**Figure 2.**
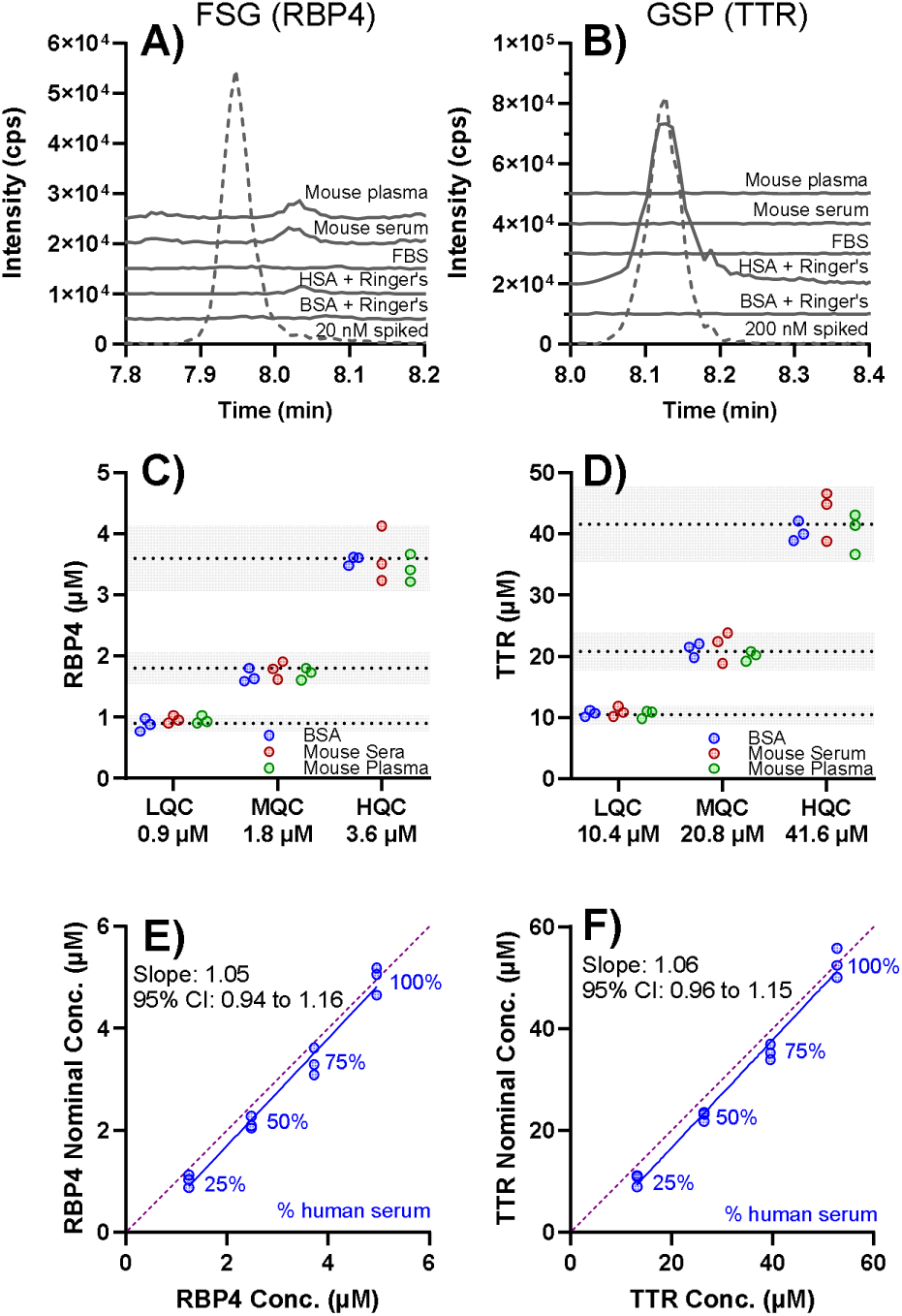
Evaluation of matrix effects, surrogate matrix, and parallelism. Chromatograms from digestions of different matrices and purified reference protein for A) RBP4 reference peptide FSG and B) TTR reference peptide GSP are shown. Solid line chromatograms depict matrix alone and dashed line chromatograms depict purified protein spiked in. Concentrations of C) RBP4 and D) TTR calculated from BSA+1% Ringers (blue), mouse serum (red), or mouse plasma (green) matrices spiked at three different concentrations of RBP4 or TTR corresponding to the low-, middle-, and high-quality controls (QC). Concentrations were quantified against a curve made in BSA+1% Ringers. The shaded area represents 15% above or below the nominal concentration. Digestions were conducted in triplicate across three separate days, results from one representative day are shown. Assessment of parallelism for E) RBP4 and F) TTR quantitation by diluting human serum into mouse serum with triplicate dilutions at each point. Each point is annotated with percent human serum. The nominal concentrations were calculated from the mean concentration of 100% human serum and adjusted based on dilution factor. The 100% mouse serum sample had no detectable peaks for either RBP4 or TTR surrogate peptides. Slope and 95% confidence intervals are inset; line of unity is a purple dashed line. BSA, bovine serum albumin; HSA, human serum albumin; FBS, fetal bovine serum.

The method was validated according to the FDA guidance and representative chromatograms are shown in **Figure 3**. Representative standard curves with QCs for RBP4 and TTR are shown in **SI Figure S6C,D**. Across 96 parallel digestions, FSG and GSP stable isotope labeled peptide peak area, along with enolase AAD peptide peak area had % CV ranging from 11 to 16% (**SI Figure S5E-H**) and the signal was stable for the analysis of the sequence.

**Figure 3.**
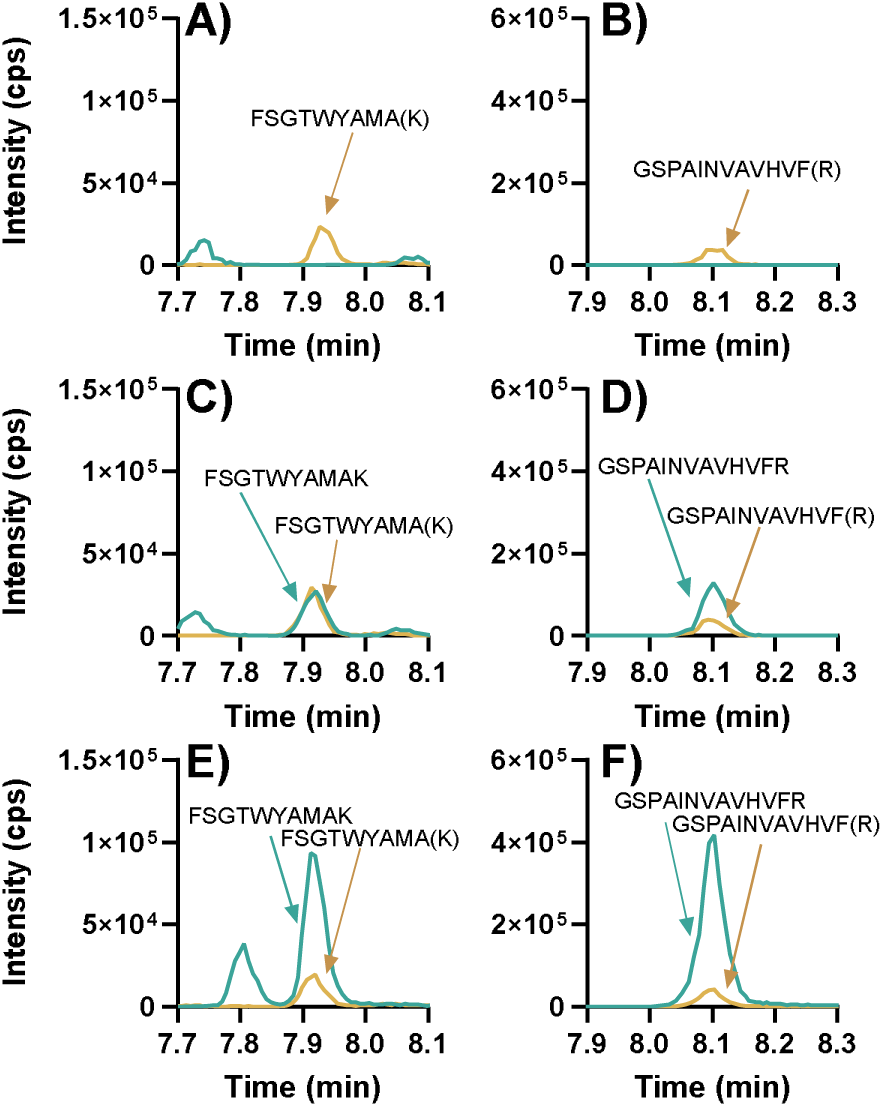
Representative chromatograms. The isotope labeled internal standard peptides for RBP4 peptide FSG (A, C, E) and for TTR peptide GSP (B, D, F) are in yellow and the corresponding analyte peptides are in turquoise. Blank matrix with isotope labeled internal standard (A, B), spiked samples at lower limit of quantitation corresponding to 0.6 µM RBP4 and 6.9 µM TTR (C, D), and representative serum sample from a healthy individual analyzed using the final validated method (E, F). Matrix was 0.6 mg/mL bovine serum albumin with 1% Ringer’s solution. Chromatograms are the sum of the three transitions used for quantification summarized in **SI Table S4**. Each panel lists the peptide sequence detected and internal standard peptide with the labelled amino acid (K[^13^C ^15^N] or R[^13^C ^15^N]) in parentheses.

At LLOQ the signal-to-noise was >>10, and peaks for the FSG and GSP peptides in a representative serum sample were greater than LLOQ across test samples (**Figure 3**). Carryover was assessed based on detection of a peak in an acetonitrile injection after the highest standard. No carryover was observed for either RBP4 or TTR. The intra-day variance for QC samples was <10% at all concentrations and the inter-day variance was <12% including the pooled QC (**Table 1**). Digested peptides for both RBP4 and TTR maintained <15% error at all QC concentrations following 24 h at 4°C or two freeze-thaw cycles (**Table 1**).

**Table 1.** Method validation data. Accuracy (% error) and precision (% CV) of RBP4 and TTR for quality control (QC) samples across nine separate days. The observed percent error (accuracy) was calculated as (nominal conc. – observed conc.)/nominal conc. for minimum n=3 at each QC concentration. Nominal concentration refers to the sample concentration before dilution with ammonium bicarbonate. LLOQ, lower limit of quantitation; LQC, low QC; MQC, middle QC; HQC, high QC.

| Peptide<br>(Protein) | QC | Nominal<br>Conc.<br>(μM) | Inter-day |  | Intra-day<br>variance<br>(%CV)<sup>a</sup> | Freeze-thaw error<br>(%)<sup>b</sup> | Autosampler<br>error (%)<sup>c</sup> |
| --- | --- | --- | --- | --- | --- | --- | --- |
|  |  |  | %<br>error | %<br>CV |  |  |  |
| **FSG**<br>**(RBP4)** | LLOQ | 0.6 | 1.5 | 11 | 5.6 | 5.6 | 11 |
|  | LQC | 0.9 | 1.3 | 8.8 | 5.9 | 2.6 | 4.1 |
|  | MQC | 1.8 | - 1.4 | 7.3 | 5.5 | 3.0 | 5.1 |
|  | HQC | 3.6 | - 4.9 | 8.7 | 5.6 | 7.5 | 9.4 |
|  | Pooled<br>QC<sup>d</sup> | – | – | 12 | 6.6 | – | – |
| **GSP**<br>**(TTR)** | LLOQ | 6.9 | - 3.8 | 9.8 | 9.8 | 9.4 | 8.1 |
|  | LQC | 10.4 | - 2.2 | 8.6 | 8.3 | 1.9 | 4.9 |
|  | MQC | 20.8 | - 4.7 | 7.6 | 6.2 | 2.8 | -1.9 |
|  | HQC | 41.6 | - 1.0 | 8.7 | 8.0 | 10.7 | -0.9 |
|  | Pooled<br>QC<sup>d</sup> | – | – | 12 | 6.6 | – | – |

<sup>a</sup>Intra-day variance was calculated from five replicate analyses within a day and the average intra-day %CV for each QC concentration from four days is reported. <sup>b</sup>Stability of quantitative peptides determined following two freeze-thaw cycles at -20°C. <sup>c</sup>Stability of quantitative peptides determined following 24 h at 4 °C in autosampler. <sup>d</sup>Pooled QC was prepared as a mixture of serum from ten individuals.

The impact of adding SIL peptides after digestion versus at the beginning of sample preparation was evaluated. Addition of SIL peptides before or after digestion resulted in comparable quantitation of RBP4 and TTR surrogate peptides, with difference less than 10% across all QC concentrations and the pooled human serum QC (**SI Table S5**).

### Method Comparison and Application

The “gold standard” method for RBP4 measurement is uncertain, particularly without reference protein standards. Historically ELISAs have been used for RBP4 quantitation; however, uncertainty in measurements has been noted in the field, particularly in samples from patients with disease or chronic conditions.^21^ In our preliminary experiments we observed similar lack of rigor in ELISA kits for RBP4 analysis. Measurement of the same serum samples using two different ELISA kits resulted in mean RBP4 concentrations that differed by 3.6-fold (data not shown). Here, RBP4 and TTR concentrations in 49 samples from individuals with diabetes and CKD were measured by ELISAs previously described in published clinical research.^12,17^ The Bland-Altman plot indicated minimal bias for both RBP4 and TTR, around 1%; however, the standard deviation of bias was greater for TTR (31%) compared to RBP4 (18%) (**Figure 4A,B**). The Pearson coefficient was 0.84 for RBP4 and 0.63 for TTR. The slope of the Deming regression was 1.05 (95% CI: 0.81-1.30) for RBP4 and 0.97 (95% CI: 0.63-1.31) for TTR, again indicating good overall agreement between the ELISAs chosen and LC-MS/MS methods but a greater variability in TTR measurements compared to RBP4 (**Figure 4C,D**). In relation to clinical use, both RBP4 and TTR measurements met the FDA incurred sample reanalysis criteria^27^ with 98% and 83% of measurements falling within 20% of the mean concentration, respectively. Geometric mean (interquartile range) concentrations measured by LC-MS/MS were 2.7 (2.4, 3.5) µM for RBP4 and 19 (17, 22) µM for TTR, in agreement with prior measurements by ELISAs in patients with CKD.^12,32^

**Figure 4.**
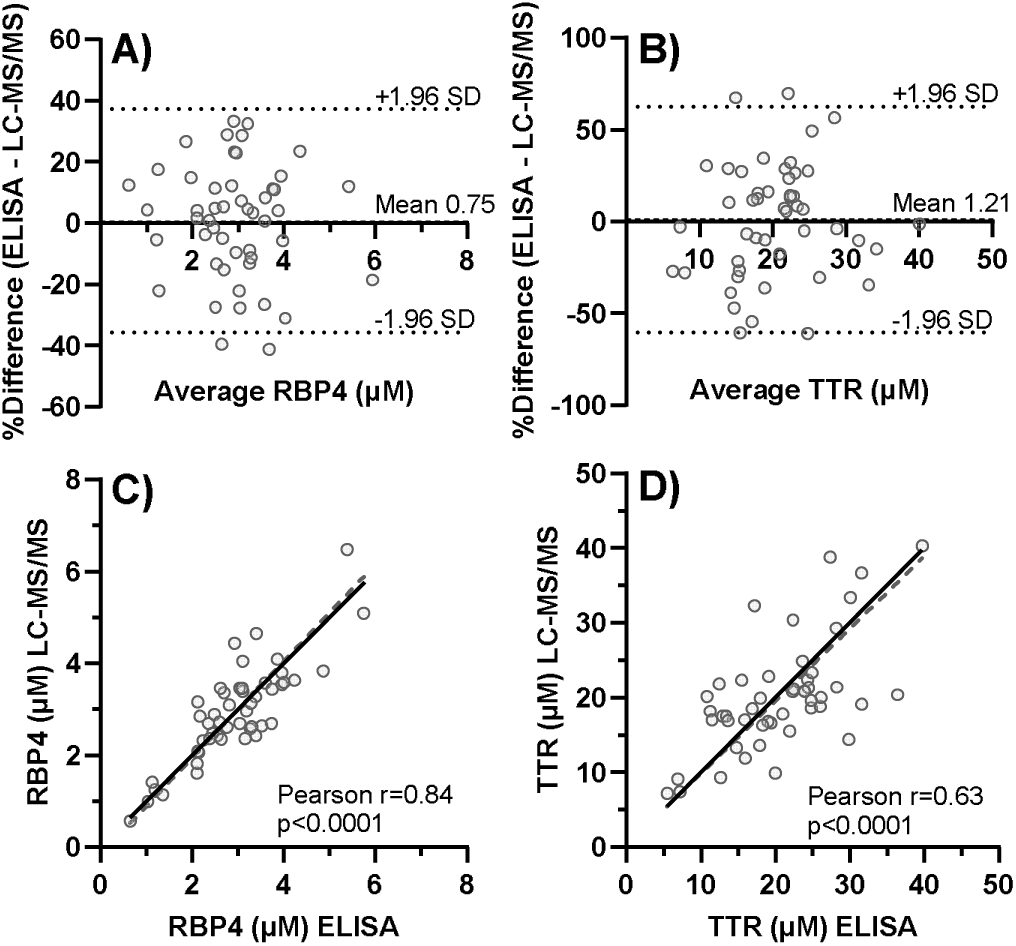
Comparison of LC-MS/MS and ELISA methods for RBP4 and TTR measurement in individuals (n=49) with chronic kidney disease (CKD). Bland-Altman plot comparing percent difference of A) RBP4 and B) TTR measurements from LC-MS/MS and ELISA. Dotted lines show the 95% confidence interval at ±1.96*standard deviation of bias, the dashed line represents mean percent bias. Deming regression for C) RBP4 ([LC-MS/MS] = 1.05*[ELISA]-0.16) and D) TTR ([LC-MS/MS]=0.94*[ELISA]+0.78) with the regression shown as a dashed line and equivalence as a solid black line. The Pearson r and p-value are inset in the figure.

TTR and RBP4 were also quantified in plasma and serum samples collected on two occasions from healthy women (n=12). The measured concentrations in matched plasma and serum samples were within 15% of each other and the average TTR:RBP4 ratio was 12 (**Table 2**). Formation of a complex between RBP4 and tetrameric TTR in circulation results in a protein complex too large for glomerular filtration,^1,3^ underscoring the critical role of TTR in conserving circulating RBP4. Variations in the TTR:RBP4 ratio may reflect physiological or pathological changes that may alter half-life of total RBP4 in circulation. The inter-individual variance for RBP4 and TTR was 18 – 26% in the healthy individuals (**Table 2**). The intra-individual variability of the TTR:RBP4 ratio was about half of this. The observed apparent intra-individual variability was similar in magnitude as the assay intra-day variance, suggesting minimal variability in RBP4 and TTR concentrations from day to day within an individual and reflecting the tight control of RBP4 and TTR in circulation (**Table 2**).

**Table 2.** Measured serum and plasma concentrations and inter- and intra-individual variability of RBP4, TTR, and TTR:RBP4 ratio in healthy female volunteers.

| Analyte | Measured concentration<br>geometric mean (range) |  | Inter-individual<br>variability (%CV) |  | Intra-individual<br>variability % |  |
| --- | --- | --- | --- | --- | --- | --- |
|  | *Serum* | *Plasma* | *Serum* | *Plasma* | *Serum* | *Plasma* |
| **RBP4<sup>a</sup>** | 1.5 (0.9 – 2.4) μM | 1.3 (1.0 – 2.1) μM | 26% | 19% | 8.1% | 8.5% |
| **TTR<sup>a</sup>** | 19 (13 – 29) μM | 16 (12 – 23) μM | 21% | 18% | 7.6% | 6.1% |
| **TTR:RBP4** | 12 (9.6 – 18) | 12 (9.1 – 15) | 13% | 9.4% | 6.3% | 5.9% |

<sup>a</sup>The RBP4 and TTR concentration in one sample was identified as an outlier using Grubbs' test ( $\alpha=0.01$ ) and excluded from the analysis.

## DISCUSSION

Quantifying RBP4 may lend mechanistic insight into physiological changes associated with numerous clinical conditions or provide a biomarker for disease progression.^4,10,11,13,14^ However, RBP4 forms a complex with TTR, and the complex may be different from free RBP4 both functionally *in vivo* and bioanalytically. Importantly, it is not known if RBP4-TTR complex formation impacts the quantitation of RBP4 or TTR by ELISAs. It is likely different ELISA kits have different limitations for quantitation. The use of different methods for RBP4 measurement may be a major source of discordance in clinical studies.^11,21^ Disease and physiologic dysregulation may similarly confound measurement of TTR by ELISA. Quantitative proteomics offers a potential solution to these confounders, and allows independent quantitation of samples to verify and cross-validate ELISA assays. Plasma proteomic analysis can, however, be affected by aspects of human plasma such as high abundance of albumin and IgG, and the presence of protease inhibitors which can affect tryptic digestion.^33,34^ Additionally, since RBP4 and TTR are endogenous proteins, the lack of a blank matrix can be problematic. We developed and validated an LC-MS/MS based method for the quantitation of RBP4 and TTR in plasma and serum that addresses these challenges. The plasma was diluted sufficiently to avoid effects of protease inhibitors and to minimize impact of albumin and IgG. Surrogate matrices that mimic human plasma were developed and evaluated allowing for preparation of standard curves and method validation. The developed method was applied to quantify RBP4 and TTR in both a healthy population and patients with diabetic CKD.

The use of the purified proteins as calibration standards and SIL peptides as internal standards allowed characterization of digestion and detection efficiency from different matrices and yielded confidence in the absolute quantitation based on the surrogate peptides. The selection and optimization of quantitative peptides in our method development emphasizes the importance of addressing digestion conditions and the need for rigorous assessment of quantitative peptides when limited options are available. For example, TTR missed cleavage peptide behaved as expected with decreasing signal as trypsin digestion time increased but the RBP4 peptide MKY in serum did not. This may be due to protease inhibitors in the matrix, that inhibit trypsin activity and lead to increased missed cleavages.^33^ Yet, the missed cleavage peptide was observed in purified protein, a matrix free of protease inhibitors, regardless of purified protein source or trypsin digest time. The source or expression system of purified RBP4 did not affect the trypsin digestion efficiency, likely because RBP4 has limited annotated post-translational modifications.^10^ The impact of detergent on protein digestion and surrogate peptide detection was highly protein specific. Sodium deoxycholate was imperative for TTR peptide signal but had minimal impact on RBP4 peptide detection. The necessity of detergent has been observed previously with a different TTR peptide.^35^

The accuracy and precision of the final method met validation criteria both for spiked samples and the pooled QC. The pooled QC sample served as a measure of assay reproducibility in real world samples. Instrument variance may account for over 50% of assay variability observed within a day. The intra-day assay variance ranged from 5.5-9.8% whereas the injection variance for a single sample ranged from 3.5-5.2%. Overall assay variance is likely a major contributor to the observed apparent intra-individual variability in RBP4 and TTR concentrations. The lowest intra-individual variance in RBP4 and TTR concentrations is the intra-day variance defined by the pooled QC. The magnitude of observed intra-individual variability in RBP4 and TTR concentrations in plasma and serum was 6-8.5%, about the same magnitude as the assay variance. Hence it is difficult to differentiate biological intra-individual variability from inherent method and instrument variance underscoring the minimal variability and homeostatic control of circulating RBP4 and TTR concentrations. The inter-individual variability was about 2-fold greater than the assay’s intra-day variance.

A challenge in developing LC-MS/MS based quantitation methods for RBP4 and TTR is their relatively short sequences. RBP4 and TTR are comprised of 183 and 147 amino acids respectively, resulting in a limited number of tryptic peptides. Indeed, only two predicted RBP4 peptides met ideal surrogate peptide selection criteria, lacking methionine, cysteine, or ragged ends, but only one of these (YWG) was experimentally observed. However, a missed cleavage (MKY) was observed in digestions of RBP4 for that peptide and thus a different peptide (FSG), despite the presence of a methionine residue, was selected for quantification. Oxidation on the methionine in the FSG peptide was not observed when tested. Similar surrogate peptide selection was previously reported for RBP4.^36^ While methionine oxidation of the FSG peptide was not observed during digestion, it remains possible that oxidation may occur prior to digestion during sample handling or storage. Thus, methionine oxidation should still be assessed, particularly when analyzing archived samples.

A limitation for all RBP4 and TTR measurement methods is the lack of certified reference standards and reference methods. This prevents confirmation of accurate measurements from relevant biological samples. Here we used multiple sources of reference RBP4 to address this issue and compared the quantitation methods to previously used ELISA assays to define accuracy of the developed method. While the results were in overall agreement with minimal bias, variance of measurements was 31% and 18% for TTR and RBP4, respectively. Based on the validation and measured assay variance of the LC-MS/MS method, the deviation is likely derived from the ELISA measurement rather than the LC-MS/MS. The shortcomings of ELISA are well-characterized,^20^ and measurements are prone to variability, as demonstrated by a 3.6-fold difference in RBP4 concentrations observed from the same serum samples analyzed with two different ELISA kits. Although our reference standard concentrations were verified using orthogonal approaches, laboratories that may apply the developed method should independently confirm the concentration of purified proteins standards. For clinical implementation, amino acid analysis together with BCA and purity assessment is likely necessary to ensure measurement accuracy.

The intra-individual variance in RBP4 and TTR concentrations was determined here from two separate visits and in a population limited to reproductive age women. This somewhat limits the generalizability of intra-individual variance findings. However, establishing baseline variability in relatively homogeneous population is a critical first step toward understanding intra-individual variance. Our findings are in agreement with a prior analysis of repeated RBP4 ELISA measurements, in which intra-individual variance was not different between men and women.^37^ Application of our assay in a clinical population comprising of both male and female patients with diabetic kidney disease demonstrates its relevance and utility in a clinical setting.

In conclusion, our method is robust and reliable for absolute quantitation of RBP4 and TTR in both serum and plasma. LC-MS/MS and ELISA measurements were overall comparable in samples from patients with diabetic CKD, though there was a larger spread in TTR measurements. It is possible this is due to sample complexity as increased variance in RBP4 measurement has been previously noted in individuals with diabetes compared to healthy volunteers.^21^ Our LC-MS/MS method offers a robust alternative to ELISAs, which have been shown to be affected by protein conformational diversity, disease states, and lack of concordance.^20^ Here, the specific peptides used for quantitation are known and the specificity of the method has been tested in multiple matrices. Our method is transferable and represents a candidate reference method for the standardization of RBP4 and TTR measurements, a critical step for the use of RBP4 as a biomarker. Our method allows for measurement of absolute concentrations of both RBP4 and TTR, enabling assessment of the stoichiometry and binding equilibria of RBP4, TTR, and retinol in circulation. This can provide insights into the proportion of total circulating RBP4 bound to retinol or complexed with TTR in serum, and how these associations may shift in clinical populations in cross-sectional or longitudinal studies. Distinction between retinol-bound and unbound RBP4 addresses a key unresolved question in the use of RBP4 as a cardiometabolic biomarker.

## ASSOCIATED CONTENT

### Supporting Information

The supporting information is available free of charge.

Supplemental Tables and Figures showing Results for RBP4 and TTR method development (DOC).

**Table S1.** Details of purified purchased RBP4 proteins

**Table S2.** Amino acid analysis of purified RBP4 and TTR reference solutions

**Table S3.** Multiple reaction monitoring (MRM) parameters, and monitored precursor and fragment ions for detected tryptic and single missed cleavage peptides of RBP4 and TTR

**Table S4.** Multiple reaction monitoring (MRM) parameters, and monitored precursor and fragment ions for quantitative peptides of RBP4, TTR, and enolase, along with stable isotope-labeled peptides for RBP4 and TTR

**Table S5.** Difference in measured concentrations of RBP4 and TTR QCs based on timing of SIL peptide addition

**Figure S1.** Detection of RBP4 peptides from initial peptide screen including initial y or b-ions and MS/MS transitions monitored

**Figure S2.** Detection of TTR peptides from initial peptide screen including initial y or b-ions and MS/MS transitions monitored.

**Figure S3.** RBP4 potential surrogate peptide peak area from trypsin digest time course of RBP4 from multiple sources

**Figure S4.** Evaluation of oxidation on methionine in FSGTWYAMAK peptide in a digested standard

**Figure S5.** Detergent effect on RBP4 and TTR peptide detection in serum and purified protein digestions

**Figure S6.** Peptide signal response linearity, standard curve and quality controls, stable isotope labeled peptide peak area, and process control enolase peptide peak area across a digestion batch

## AUTHOR INFORMATION

### Author Contributions

A.S.Y., A.Z., and N.I. designed the experiments. A.S.Y. performed experiments and data analysis. K.B.R., A.K.A., J.K.A., and N.I. designed and oversaw the healthy volunteer clinical study and provided the samples analyzed from healthy volunteers. B.R.K. provided the samples of the kidney disease study. N.I. and K.B.R acquired funding for the project leading to this publication. A.S.Y and N.I. wrote the manuscript, all authors reviewed and edited the manuscript.

The manuscript was written through contributions of all authors. All authors have given approval to the final version of the manuscript.

### Notes

The authors declare no competing financial interest

## ACKNOWLEDGMENT

The work presented here was supported by National Institutes of Health Grants R01-GM111772 [ASY, KBR, NI, AZ], P01-DA032507 [KBR, AKA, JKA, NI, AZ], T32-GM007750 [ASY, AKA], TL1-TR002318 [ASY]. Support was also provided by the UW School of Pharmacy’s Milo Gibaldi Endowed Chair [NI] and William E. Bradley Endowed Fellowship [ASY].

We thank Drs. Andrew N. Hoofnagle and Huu-Hien Huynh in the Department of Laboratory Medicine & Pathology at the University of Washington associated with the Diabetes Research Center for performing the amino acid analysis on RBP4 and TTR reference solutions. The Diabetes Research Center is funded by the National Institutes of Health P30 DK07047. We thank the Translational Research Unit, which is a part of Institute of Translational Health Sciences funded by the National Institutes of Health UL1TR002319. The TOC graphic was created in BioRender. Yadav, A. (2025) https://BioRender.com/v02q389.

## ABBREVIATIONS

RBP4: retinol binding protein 4
TTR: transthyretin
ELISA: enzyme linked immunoassay
LC-MS/MS: liquid chromatograph-tandem mass spectrometry
SIL: stable isotope labeled
HSA: human serum albumin
BSA: bovine serum albumin
FBS: fetal bovine serum
DTT: dithiothreitol
TCEP: tris(2-carboxyethyl)phosphine
TFA: trifluoroacetic acid
QC: quality control
LLOQ: lower limit of quantitation
LQC: low QC
MQC: medium QC
HQC: high QC
CKD: chronic kidney disease
IQR: interquartile range

---

For Table of Contents Only

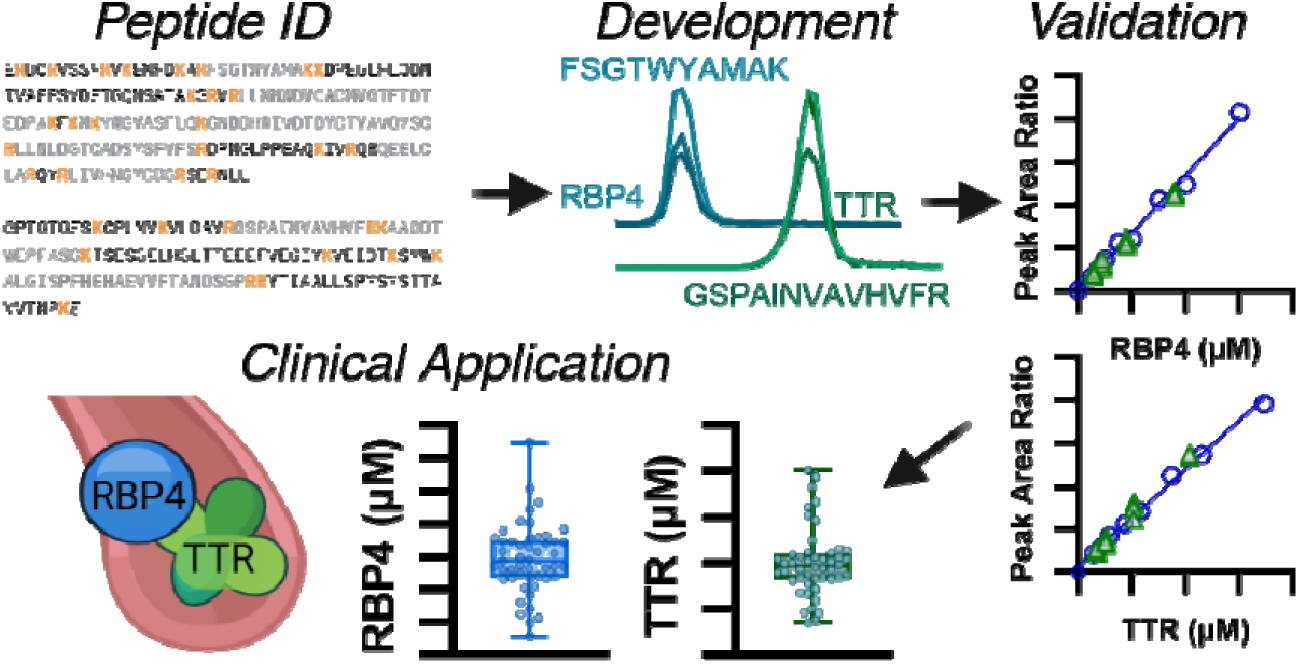

## REFERENCES

(1) Kanai, M.; Raz, A.; Goodman, D. S. Retinol-Binding Protein: The Transport Protein for Vitamin A in Human Plasma. J. Clin. Invest. 1968, 47 (9), 2025–2044. 10.1172/JCI105889.

(2) Goodman, D. S. Plasma Retinol-Binding Protein. In The Retinoids; Elsevier, 1984; pp 41–88. 10.1016/B978-0-12-658102-7.50008-7.

(3) Van Bennekum, A. M.; Wei, S.; Gamble, M. V.; Vogel, S.; Piantedosi, R.; Gottesman, M.; Episkopou, V.; Blaner, W. S. Biochemical Basis for Depressed Serum Retinol Levels in Transthyretin-Deficient Mice. J Biol Chem 2001, 276 (2), 1107–1113. 10.1074/jbc.M008091200.

(4) Yadav, A. S.; Isoherranen, N.; Rubinow, K. B. Vitamin A Homeostasis and Cardiometabolic Disease in Humans: Lost in Translation? Journal of Molecular Endocrinology 2022, 69 (3), R95–R108. 10.1530/JME-22-0078.

(5) Ingenbleek, Y.; Young, V. Transthyretin (Prealbumin) in Health and Disease: Nutritional Implications. Annu. Rev. Nutr. 1994, 14 (1), 495–533. 10.1146/annurev.nu.14.070194.002431.

(6) Underwood, B. A. Vitamin A in Animal and Human Nutrition. In The Retinoids; Elsevier, 1984; pp 281–392. 10.1016/B978-0-12-658101-0.50012-3.

(7) Shirakami, Y.; Lee, S.-A.; Clugston, R. D.; Blaner, W. S. Hepatic Metabolism of Retinoids and Disease Associations. Biochim Biophys Acta 2012, 1821 (1), 124–136. 10.1016/j.bbalip.2011.06.023.

(8) Muto, Y.; Smith, J. E.; Milch, P. O.; Goodman, D. S. Regulation of Retinol-Binding Protein Metabolism by Vitamin A Status in the Rat. J Biol Chem 1972, 247 (8), 2542–2550. 10.1016/S0021-9258(19)45460-4.

(9) Baeten, J. M.; Richardson, B. A.; Bankson, D. D.; Wener, M. H.; Kreiss, J. K.; Lavreys, L.; Mandaliya, K.; Bwayo, J. J.; McClelland, R. S. Use of Serum Retinol-Binding Protein for Prediction of Vitamin A Deficiency: Effects of HIV-1 Infection, Protein Malnutrition, and the Acute Phase Response. Am J Clin Nutr 2004, 79 (2), 218–225. 10.1093/ajcn/79.2.218.

(10) Steinhoff, J. S.; Lass, A.; Schupp, M. Biological Functions of RBP4 and Its Relevance for Human Diseases. Front. Physiol. 2021, 12, 659977. 10.3389/fphys.2021.659977.

(11) Blaner, W. S. Vitamin A Signaling and Homeostasis in Obesity, Diabetes, and Metabolic Disorders. Pharmacol Ther 2019, 197, 153–178. 10.1016/j.pharmthera.2019.01.006.

(12) Jing, J.; Isoherranen, N.; Robinson-Cohen, C.; Petrie, I.; Kestenbaum, B. R.; Yeung, C. K. Chronic Kidney Disease Alters Vitamin A Homeostasis via Effects on Hepatic RBP4 Protein Expression and Metabolic Enzymes. Clin Transl Sci 2016, 9 (4), 207–215. 10.1111/cts.12402.

(13) Graham, T. E.; Yang, Q.; Blüher, M.; Hammarstedt, A.; Ciaraldi, T. P.; Henry, R. R.; Wason, C. J.; Oberbach, A.; Jansson, P.-A.; Smith, U.; Kahn, B. B. Retinol-Binding Protein 4 and Insulin Resistance in Lean, Obese, and Diabetic Subjects. N Engl J Med 2006, 354 (24), 2552–2563. 10.1056/NEJMoa054862.

(14) Sun, Q.; Kiernan, U. A.; Shi, L.; Phillips, D. A.; Kahn, B. B.; Hu, F. B.; Manson, J. E.; Albert, C. M.; Rexrode, K. M. Plasma Retinol-Binding Protein 4 (RBP4) Levels and Risk of Coronary Heart Disease: A Prospective Analysis Among Women in the Nurses’ Health Study. Circulation 2013, 127 (19), 1938–1947. 10.1161/CIRCULATIONAHA.113.002073.

(15) Leca, B. M.; Kite, C.; Lagojda, L.; Davasgaium, A.; Dallaway, A.; Chatha, K. K.; Randeva, H. S.; Kyrou, I. Retinol-Binding Protein 4 (RBP4) Circulating Levels and Gestational Diabetes Mellitus: A Systematic Review and Meta-Analysis. Front. Public Health 2024, 12, 1348970. 10.3389/fpubh.2024.1348970.

(16) Hamdan, H. Z.; Ali, T.; Adam, I. Association between Retinol-Binding Protein 4 Levels and Preeclampsia: A Systematic Review and Meta-Analysis. Nutrients 2022, 14 (24), 5201. 10.3390/nu14245201.

(17) Czuba, L. C.; Fay, E. E.; LaFrance, J.; Smith, C. K.; Shum, S.; Moreni, S. L.; Mao, J.; Isoherranen, N.; Hebert, M. F. Plasma Retinoid Concentrations Are Altered in Pregnant Women. Nutrients 2022, 14 (7), 1365. 10.3390/nu14071365.

(18) Vilà-Rico, M.; Colomé-Calls, N.; Martín-Castel, L.; Gay, M.; Azorín, S.; Vilaseca, M.; Planas, A.; Canals, F. Quantitative Analysis of Post-Translational Modifications in Human Serum Transthyretin Associated with Familial Amyloidotic Polyneuropathy by Targeted LC–MS and Intact Protein MS. J. Proteomics 2015, 127, 234–246. 10.1016/j.jprot.2015.04.016.

(19) Poulsen, K.; Bahl, J. M. C.; Tanassi, J. T.; Simonsen, A. H.; Heegaard, N. H. H. Characterization and Stability of Transthyretin Isoforms in Cerebrospinal Fluid Examined by Immunoprecipitation and High-Resolution Mass Spectrometry of Intact Protein. Methods 2012, 56 (2), 284–292. 10.1016/j.ymeth.2011.12.009.

(20) Hoofnagle, A. N.; Wener, M. H. The Fundamental Flaws of Immunoassays and Potential Solutions Using Tandem Mass Spectrometry. J Immunol Methods 2009, 347 (1–2), 3–11. 10.1016/j.jim.2009.06.003.

(21) Graham, T. E.; Wason, C. J.; Blüher, M.; Kahn, B. B. Shortcomings in Methodology Complicate Measurements of Serum Retinol Binding Protein (RBP4) in Insulin-Resistant Human Subjects. Diabetologia 2007, 50 (4), 814–823. 10.1007/s00125-006-0557-0.

(22) Amory, J.; Muller, C.; Walsh, T. Isotretinoin for the Treatment of Nonobstructive Azoospermia: A Pilot Study. Asian J Androl 2021, 23 (5), 537. 10.4103/aja.aja_18_21.

(23) Gasteiger, E.; Hoogland, C.; Gattiker, A.; Duvaud, S.; Wilkins, M. R.; Appel, R. D.; Bairoch, A. Protein Identification and Analysis Tools on the ExPASy Server. In The Proteomics Protocols Handbook; Walker, J. M., Ed.; Humana Press: Totowa, NJ, 2005; pp 571–607. 10.1385/1-59259-890-0:571.

(24) Huynh, H.-H.; Kuch, K.; Orquillas, A.; Forrest, K.; Barahona-Carrillo, L.; Keene, D.; Henderson, V. W.; Wagner, A. D.; Poston, K. L.; Montine, T. J.; Lin, A.; Tian, L.; MacCoss, M. J.; Emrick, M. A.; Hoofnagle, A. N. Metrologically Traceable Quantification of 3 Apolipoprotein E Isoforms in Cerebrospinal Fluid. Clinical Chemistry 2023, 69 (7), 734–745. 10.1093/clinchem/hvad056.

(25) Pino, L. K.; Searle, B. C.; Bollinger, J. G.; Nunn, B.; MacLean, B.; MacCoss, M. J. The Skyline Ecosystem: Informatics for Quantitative Mass Spectrometry Proteomics. Mass Spectrom Rev 2020, 39 (3), 229–244. 10.1002/mas.21540.

(26) Arnold, S. L.; Stevison, F.; Isoherranen, N. Impact of Sample Matrix on Accuracy of Peptide Quantification: Assessment of Calibrator and Internal Standard Selection and Method Validation. Anal. Chem. 2016, 88 (1), 746–753. 10.1021/acs.analchem.5b03004.

(27) U.S. Food and Drug Administration. Bioanalytical Method Validation. Guidance for Industry. 2018.

(28) Tsantilas, K. A.; Merrihew, G. E.; Robbins, J. E.; Johnson, R. S.; Park, J.; Plubell, D. L.; Canterbury, J. D.; Huang, E.; Riffle, M.; Sharma, V.; MacLean, B. X.; Eckels, J.; Wu, C. C.; Bereman, M. S.; Spencer, S. E.; Hoofnagle, A. N.; MacCoss, M. J. A Framework for Quality Control in Quantitative Proteomics. J. Proteome Res. 2024, 23 (10), 4392–4408. 10.1021/acs.jproteome.4c00363.

(29) Bosworth, C. R.; Levin, G.; Robinson-Cohen, C.; Hoofnagle, A. N.; Ruzinski, J.; Young, B.; Schwartz, S. M.; Himmelfarb, J.; Kestenbaum, B.; de Boer, I. H. The Serum 24,25-Dihydroxyvitamin D Concentration, a Marker of Vitamin D Catabolism, Is Reduced in Chronic Kidney Disease. Kidney Int. 2012, 82 (6), 693–700. 10.1038/ki.2012.193.

(30) Sharma, V.; Eckels, J.; Schilling, B.; Ludwig, C.; Jaffe, J. D.; MacCoss, M. J.; MacLean, B. Panorama Public: A Public Repository for Quantitative Data Sets Processed in Skyline. Mol Cell Proteomics. 2018, 17 (6), 1239–1244. 10.1074/mcp.RA117.000543.

(31) Pino, L. K.; Searle, B. C.; Yang, H.-Y.; Hoofnagle, A. N.; Noble, W. S.; MacCoss, M. J. Matrix-Matched Calibration Curves for Assessing Analytical Figures of Merit in Quantitative Proteomics. J. Proteome Res. 2020, 19 (3), 1147–1153. 10.1021/acs.jproteome.9b00666.

(32) Frey, S. K.; Nagl, B.; Henze, A.; Raila, J.; Schlosser, B.; Berg, T.; Tepel, M.; Zidek, W.; Weickert, M. O.; Pfeiffer, A. F.; Schweigert, F. J. Isoforms of Retinol Binding Protein 4 (RBP4) Are Increased in Chronic Diseases of the Kidney but Not of the Liver. Lipids Health Dis 2008, 7 (1), 29. 10.1186/1476-511X-7-29.

(33) Clifton, J.; Huang, F.; Rucevic, M.; Cao, L.; Hixson, D.; Josic, D. Protease Inhibitors as Possible Pitfalls in Proteomic Analyses of Complex Biological Samples. J. Proteom 2011, 74 (7), 935–941. 10.1016/j.jprot.2011.02.010.

(34) Hortin, G. L.; Sviridov, D.; Anderson, N. L. High-Abundance Polypeptides of the Human Plasma Proteome Comprising the Top 4 Logs of Polypeptide Abundance. Clin Chem 2008, 54 (10), 1608–1616. 10.1373/clinchem.2008.108175.

(35) Proc, J. L.; Kuzyk, M. A.; Hardie, D. B.; Yang, J.; Smith, D. S.; Jackson, A. M.; Parker, C. E.; Borchers, C. H. A Quantitative Study of the Effects of Chaotropic Agents, Surfactants, and Solvents on the Digestion Efficiency of Human Plasma Proteins by Trypsin. J. Proteome Res. 2010, 9 (10), 5422–5437. 10.1021/pr100656u.

(36) Phipps, W. S.; Greene, D. N.; Pflaum, H.; Laha, T. J.; Dickerson, J. A.; Irvine, J.; Merrill, A. E.; Ranjitkar, P.; Henderson, C. M.; Hoofnagle, A. N. Small Volume Retinol Binding Protein Measurement by Liquid Chromatography-Tandem Mass Spectrometry. Clinical Biochemistry 2022, 99, 111–117. 10.1016/j.clinbiochem.2021.10.005.

(37) Wittenbecher, C.; di Giuseppe, R.; Biemann, R.; Menzel, J.; Arregui, M.; Hoffmann, J.; Aleksandrova, K.; Boeing, H.; Isermann, B.; Schulze, M. B.; Weikert, C. Reproducibility of Retinol Binding Protein 4 and Omentin-1 Measurements over a Four Months Period: A Reliability Study in a Cohort of 207 Apparently Healthy Participants. PLoS One 2015, 10 (9), e0138480. 10.1371/journal.pone.0138480.

