## Supplemental materials for "Development and Validation of a Novel LC-MS/MS Based Proteomics Method for Quantitation of Retinol Binding Protein 4 (RBP4) and Transthyretin (TTR)"

### Supplemental Tables and Figures

**Table S1.** Details of the purified RBP4 proteins purchased for method development. HEK 293, human embryonic kidney cells; PBS, phosphate buffered saline.

| Manufacturer | Source | Modifications | Provided in | Part Number |
|---|---|---|---|---|
| Sigma-Aldrich | *E. Coli* | - | PBS solution | SRP6038 |
| Sigma-Aldrich | HEK 293 | FLAG0174-tag | PBS solution | SRP6532 |
| R&D Systems | Myeloma | 6x-histidine tag | Lyophilized | 3378-LC |
| Abnova | HEK 293 | 6x-histidine tag | PBS with 25% glycerol | P5124 |
| Bio-Rad | Purified from human urine | - | Lyophilized | 7970-0504 |

**Table S2.** Amino acid analysis of RBP4 and TTR protein concentration, performed in triplicate.

| Protein | Theoretical conc. (mg/mL) | Protein conc. (mg/g) | | | | | Average | %CV | Calculated conc. (mg/mL) † | Accuracy (% Bias) |
|---|---|---|---|---|---|---|---|---|---|---|
| | | P | V | I | L | F | | | | |
| RBP4 | 0.25 | 0.293 | 0.249 | 0.266 | 0.211 | 0.222 | 0.248 | 13.4 | 0.254 | 1.4 % |
| TTR | 0.0826 | 0.083 | 0.084 | 0.086 | 0.092 | 0.087 | 0.086 | 3.9 | 0.088 | 6.9 % |

† Calculated peptide/protein concentration uses density = 1.0218 g/mL; P: proline, V: valine, I: isoleucine, L: leucine, F: phenylalanine.

**Table S3.** Multiple reaction monitoring (MRM) parameters, and monitored precursor and fragment ions for detected tryptic and single missed cleavage peptides of RBP4 and TTR.

| Protein | Peptide amino acid sequence | Precursor m/z | Fragment | m/z | Fragment | m/z | Fragment | m/z | DP (V) | CE (V) |
|---|---|---|---|---|---|---|---|---|---|---|
| RBP4 | FSGTWYAMAK (FSG) | $581.3^{+2}$ | y9 | 1014.5 | y8 | 927.4 | y6 | 769.4 | 80 | 25.0 |
| | LLNNWDV[C]ADMVGTFTDTEDPAK | $871.4^{+3}$ | y3 | 315.2 | y7 | 775.3 | y4 | 430.2 | 80 | 39.8 |
| | YWGVASFLQK (YWG) | $599.8^{+2}$ | y6 | 693.4 | b2 | 350.2 | b3 | 407.2 | 60 | 25.0 |
| | MKYWGVASFLQK (MKY) † | $729.4^{+2}$ | y8 | 849.5 | b3 | 423.2 | | | 60 | 34.7 |
| | | $486.6^{+3}$ | y6 | 693.4 | | | | | 60 | 22.0 |
| | GNDDHWIVDTDYDTYAVQYS[C]R | $898.4^{+3}$ | y6 | 812.4 | y5 | 713.3 | | | 80 | 41.1 |
| | LLNLDGT[C]ADSYSFVFSR | $1033.0^{+2}$ | b3 | 341.2 | | | | | 80 | 49.6 |
| | | $689.0^{+3}$ | y6 | 742.4 | | | | | 80 | 31.1 |
| | QEEL[C]LAR | $509.7^{+2}$ | y4 | 519.3 | y6 | 761.4 | y3 | 359.2 | 80 | 24 |
| | LIVHNGY[C]DGR | $435.2^{+3}$ | $y9^{+2}$ | 539.2 | y4 | 507.2 | y2 | 232.2 | 80 | 18.9 |
| TTR | GSPAINVAVHVFR (GSP) | $683.9^{+2}$ | $y11^{+2}$ | 611.9 | y8 | 941.5 | y6 | 728.4 | 80 | 32.5 |
| | GSPAINVAVHVFRK † | $747.9^{+2}$ | $y5^{+}$ | 686.4 | | | | | 80 | 35.6 |
| | | $499.0^{+3}$ | $y12^{+2}$ | 675.9 | $y10^{+2}$ | 591.9 | | | 80 | 21.9 |
| | AADDTWEPFASGK | $697.8^{+2}$ | y8 | 921.4 | y7 | 735.4 | y6 | 606.3 | 80 | 33.2 |
| | KAADDTWEPFASGK † | $761.9^{+2}$ | b8 | 917.4 | y9 | 1022.5 | | | 80 | 36.3 |
| | | $508.2^{+3}$ | y6 | 606.3 | | | | | 80 | 22.4 |
| | ALGISPFHEHAEVVFTANDSGPR | $817.7^{+3}$ | y9 | 964.5 | $b13^{+2}$ | 694.9 | $b14^{+2}$ | 744.4 | 80 | 37.3 |

† Peptide with missed cleavage. [C] indicates cysteine alkylation. Fragment refers to y or b-ions for each peptide precursor. Precursor charge state on the precursor m/z, y- and b- ion charge states are +1 unless specified. Collision cell exit potential (CXP) and entrance potential (EP) were 13 V and 10 V, respectively, for all transitions. DP: declustering potential, CE: collision energy.

**Table S4.** Multiple reaction monitoring (MRM) parameters, and monitored precursor and fragment ions for quantitative peptides of RBP4, TTR, and enolase along with stable isotope-labeled peptides for RBP4 and TTR.

| Protein | Peptide amino acid sequence | Precursor m/z | Fragment | m/z | Fragment | m/z | Fragment | m/z | CE (V) |
|---|---|---|---|---|---|---|---|---|---|
| RBP4 | FSGTWYAMAK (FSG) | $581.3^{+2}$ | y9 | 1014.5 | y8 | 927.4 | y6 | 769.4 | 25.0 |
| | FSGTWYAMAK[$^{13}C_6{}^{15}N_2$] | $585.3^{+2}$ | y9 | 1022.5 | y8 | 935.4 | y6 | 777.4 | 25.0 |
| TTR | GSPAINVAVHVFR (GSP) | $683.9^{+2}$ | $y11^{+2}$ | 611.9 | y8 | 941.5 | y6 | 728.4 | 32.5 |
| | GSPAINVAVHVFR[$^{13}C_6{}^{15}N_4$] | $688.9^{+2}$ | $y11^{+2}$ | 616.9 | y8 | 951.5 | y6 | 738.4 | 32.5 |
| Enolase | AADALLLK | $407.8^{+2}$ | y7 | 743.5 | y6 | 672.4 | | | 19.0 |
| | TFAEALR | $404.2^{+2}$ | y5 | 559.3 | b2 | 249.1 | | | 18.8 |

Fragment refers to y- or b- ions for each peptide precursor. Precursor charge state on the precursor m/z, y and b-ion charge states are +1 unless specified. Collision cell exit potential (CXP) , entrance potential (EP) and declustering potential (DP) were 13 V, 10 V, and 80 V, respectively, for all transitions. Collision energy (CE), CXP, declustering potential (DP) and EP were optimized using isotope-labeled peptides for FSG and GSP peptides. The CE was adjusted to 25 V from 27.5 V for FSG and no changes were made for GSP.

**Table S5.** Difference in measured concentrations of RBP4 and TTR when SIL peptides were added before digestion compared to after digestion. Accuracy (% difference or bias) was calculated as: (conc. with SIL peptide added before – conc. with SIL peptide added after)/ conc. with SIL peptide added after × 100. Each QC concentration was digested in triplicate under both conditions. Nominal concentration refers to the sample concentration before dilution with ammonium bicarbonate.

| Peptide (Protein) | QC | Nominal Conc. (μM) | % Bias |
|---|---|---|---|
| **FSG (RBP4)** | LLOQ | 0.6 | 9.6 |
| | LQC | 0.9 | 3.4 |
| | MQC | 1.8 | -7.8 |
| | HQC | 3.6 | 2.5 |
| | Pooled QC[a] | – | 0.3 |
| **GSP (TTR)** | LLOQ | 6.9 | 4.8 |
| | LQC | 10.4 | 7.4 |
| | MQC | 20.8 | 0.6 |
| | HQC | 41.6 | 5.6 |
| | Pooled QC[a] | – | 4.4 |

[a]Pooled QC was prepared as a mixture of serum from ten individuals.

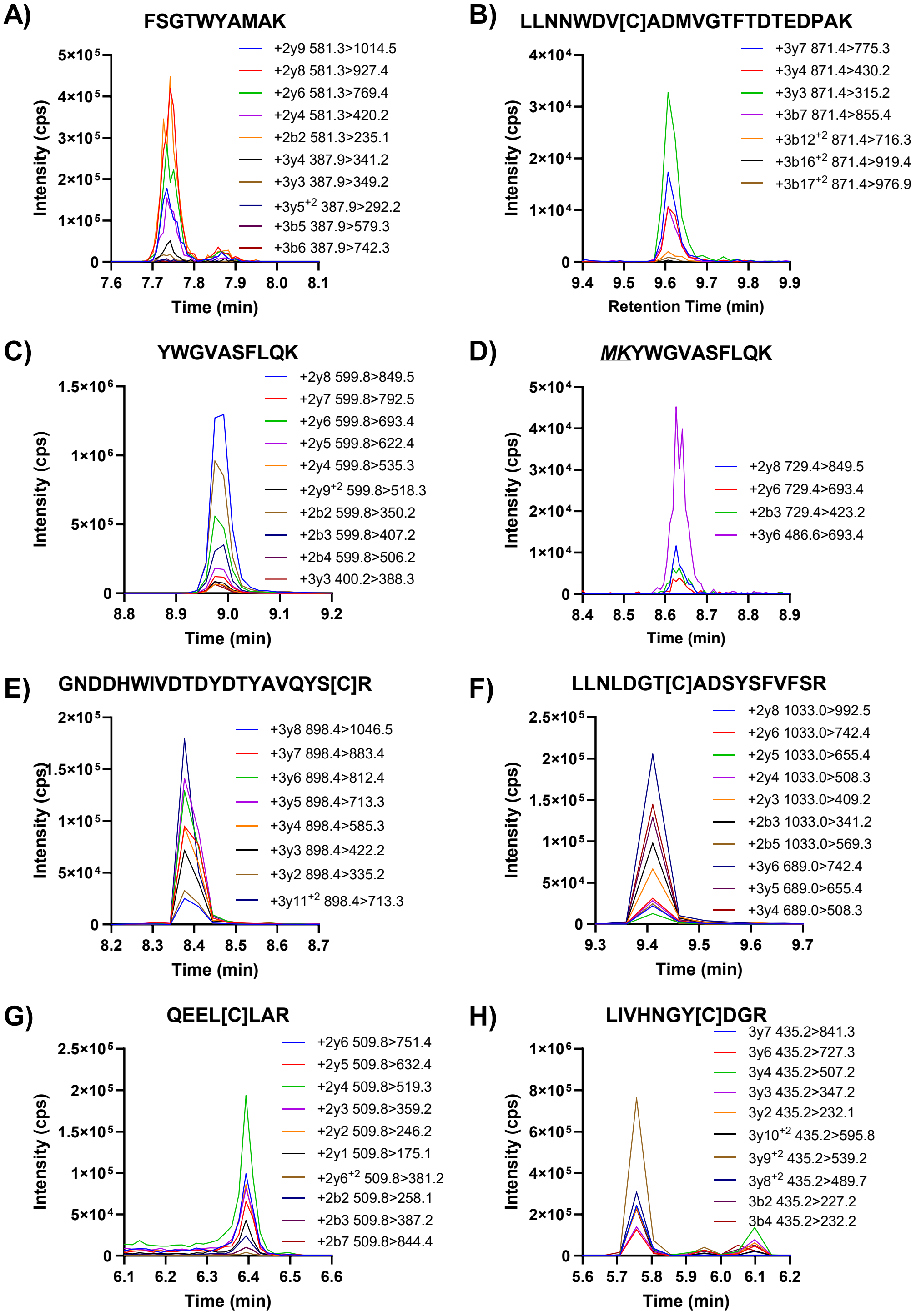

**Figure S1.** Detection of RBP4 peptides (A-H) from initial peptide screen following digestion of 400 nM purified protein. Peptides with missed cleavages (D) have the missed cleavage residues shown in italics and underlined. The alkylated cysteine residues are marked as [C]. The insets list the charge state, specific y or b-ions and the MS/MS transitions monitored and detected.

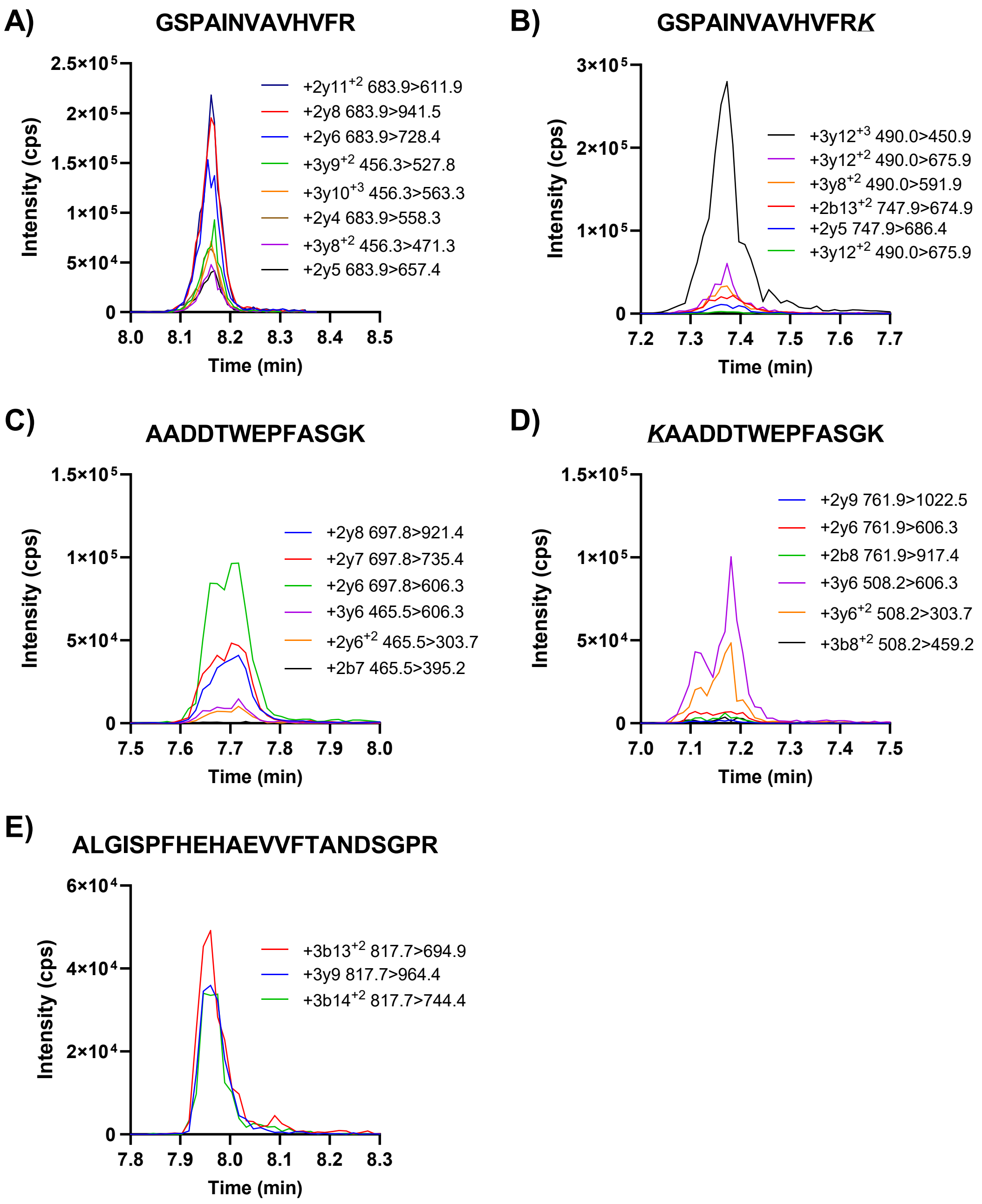

**Figure S2.** Detection of TTR peptides (A-E) from initial peptide screen following digestion of 400 nM purified protein. Peptides with missed cleavages (B,D) have the missed cleavage residues shown in italics and underlined. The insets list the charge state, specific y or b-ions and the MS/MS transitions monitored.

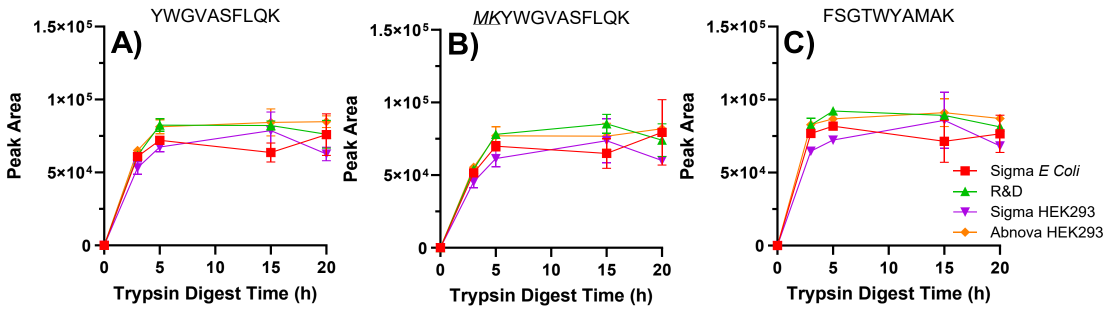

**Figure S3.** Trypsin digest time course of RBP4 from different sources of RBP4 listed in Table S1. The measured peptides shown are A) YWG, B) missed cleavage MKY, and C) FSG. Sigma E. Coli refers to RBP4 expressed in E. Coli, R&D to RBP4 expressed in mouse myeloma cells, Sigma HEK293 and Abnova HEK293 to RBP4 expressed in human embryonic kidney cells (HEK293 cells). Missed cleavage residues are underlined and italicized in B). Each digestion was carried out in triplicate.

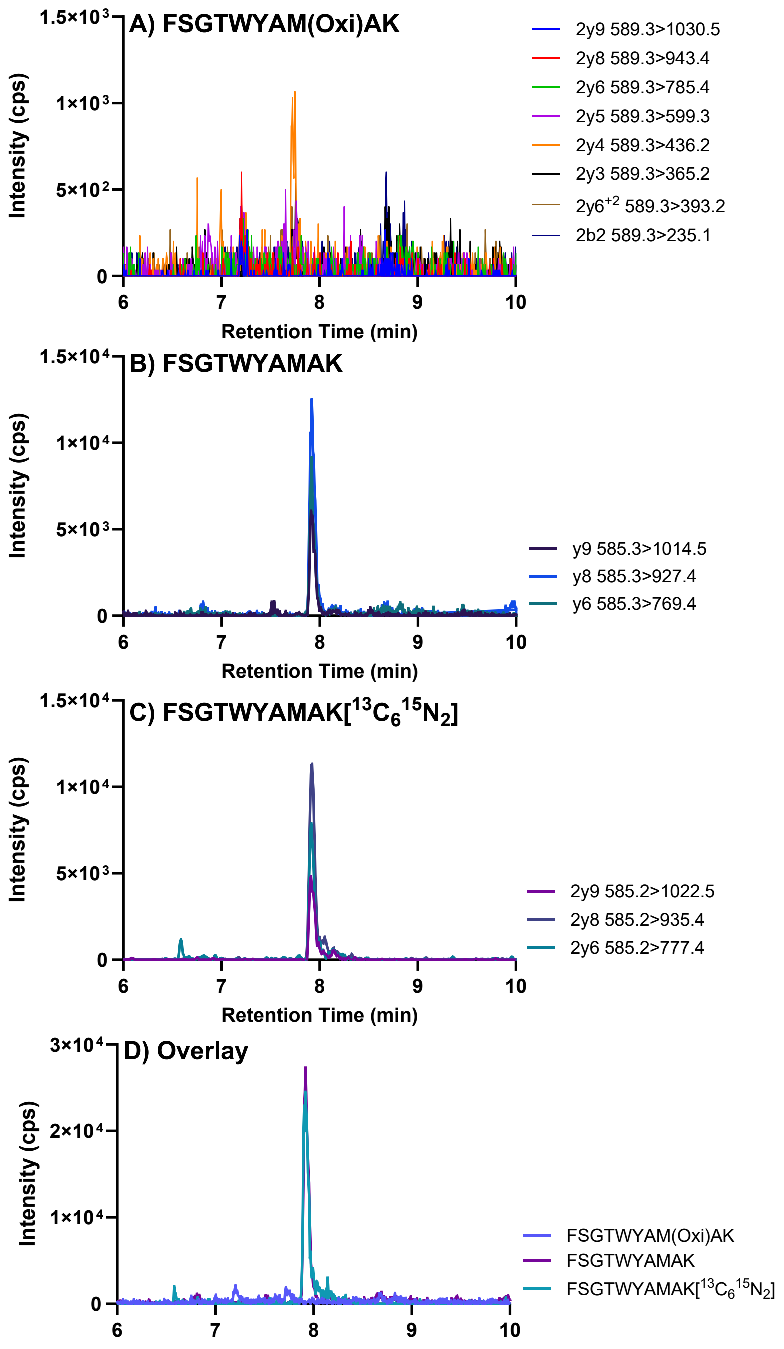

**Figure S4.** Evaluation of methionine oxidation in the FSGTWYAMAK peptide in a digested standard corresponding to 2 μM RBP4 in undiluted plasma or serum. A) Extracted ion chromatograms of select MS/MS transitions for the predicted b and y ions of the of FSGTWYAMAK peptide with methionine oxidation. B) FSG peptide response for the peptide without methionine oxidation. C) FSG stable isotope labeled peptide response. D) overlay of the chromatograms in A-C with all transitions summed for each. Insets detail the charge state, specific y and b-ions and the MS/MS transitions monitored for A through C.

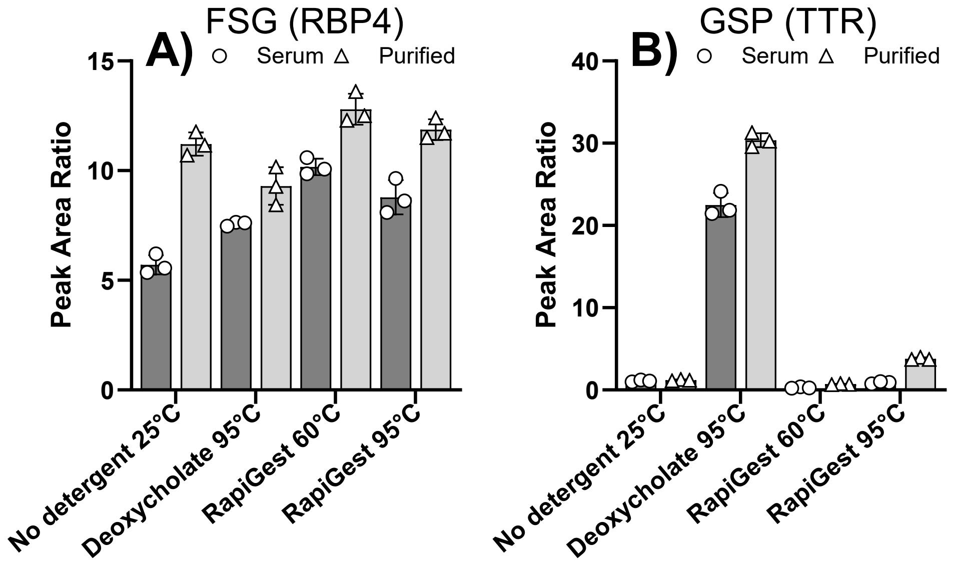

**Figure S5.** Detergent effect on RBP4 and TTR peptide detection in serum and purified protein. Peak area ratio of A) FSG and B) GSP peptides normalized to corresponding SIL peptides in serum (circles, dark gray bars) and purified protein (triangles, light gray bars). RapiGest surfactant incubations were conducted at both 60°C and 95°C. Digestions were conducted in triplicate.

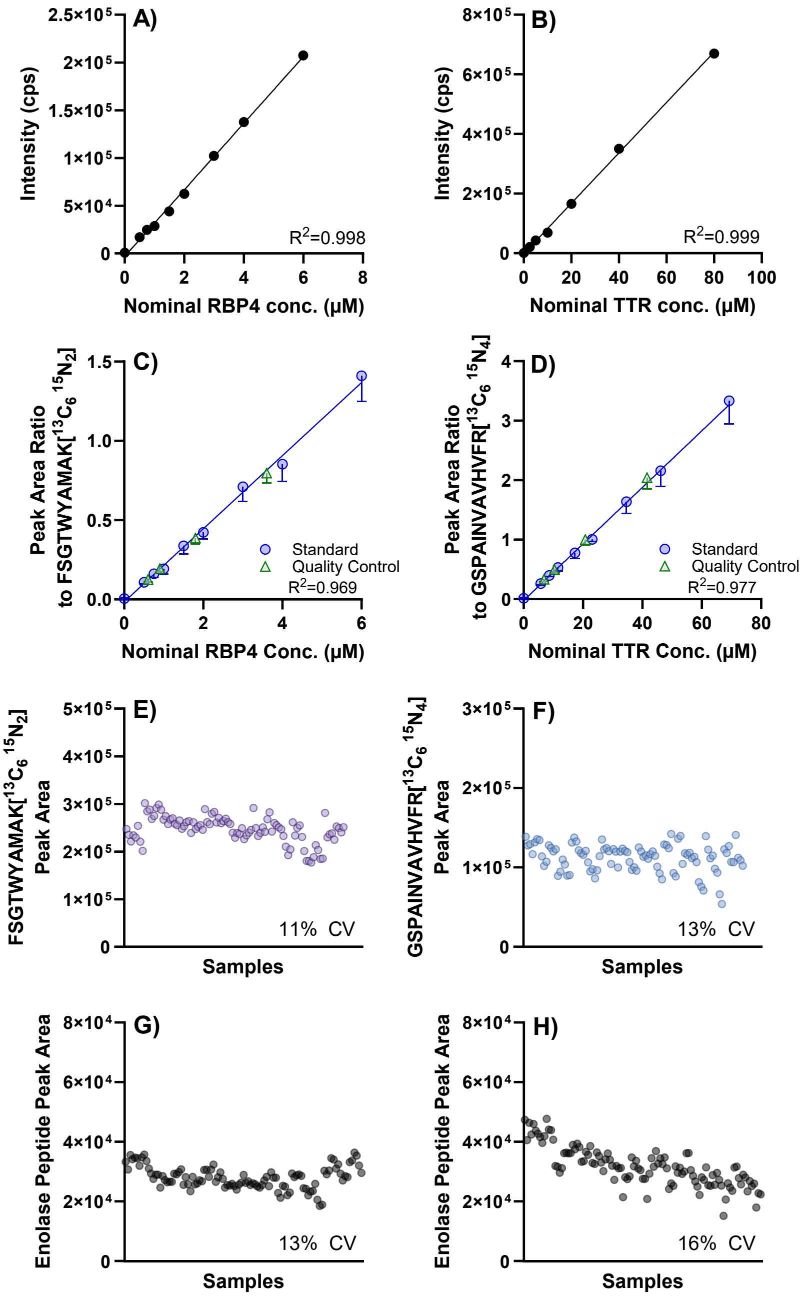

**Figure S6***.* Linearity of peptide signal response on instrument for A) FSG from 5 – 60 nM and B) GSP from 25 – 800 nM from serial dilutions (n=1), corresponding to undiluted plasma or serum concentrations of 0.5 – 6 µM RBP4 and 2.5 – 80 µM TTR. All standards were diluted 100x before digestion. The $R^2$ is from an unweighted linear regression. Mean and standard deviation of peak area ratios from six standard curves (blue circles) and 18 sets of quality control samples (green triangles) for C) RBP4 using FSG peptide and D) TTR using GSP peptide. The $R^2$ is from a linear regression with 1/x weighting. Stable isotope-labeled peptide peak area across a 96-well plate of digestions conducted in parallel for E) FSGTWYAMAK[$^{13}C_6{}^{15}N_2$] and F) GSPAINVAVHVFR[$^{13}C_6{}^{15}N_4$]. Enolase peptide AADALLLK signal response on two separate analysis days G) and H) from across 96 digestions conducted in parallel on each day. %CV: coefficient of variance.
